# SIGNR1 promotes immune dysfunction in systemic candidiasis by modulating neutrophil lifespan via T cell-derived histones and G-CSF

**DOI:** 10.1101/2021.08.09.455510

**Authors:** Marianna Ioannou, Dennis Hoving, Iker Valle Aramburu, Nathalia M. De Vasconcelos, Mia I. Temkin, Qian Wang, Spyros Vernardis, Vadim Demichev, Theodora-Dorita Tsourouktsoglou, Stefan Boeing, Robert Goldstone, Sascha David, Klaus Stahl, Christian Bode, Markus Ralser, Venizelos Papayannopoulos

**Author notes:** these authors contributed equally (Marianna Ioannou, Dennis Hoving).

## Abstract

The mechanisms regulating immune dysfunction during sepsis are poorly understood. Here, we show that neutrophil-derived myeloperoxidase delays the onset of immune dysfunction during systemic candidiasis by controlling microbes captured by splenic marginal zone (MZ) macrophages. In contrast, SIGNR1-mediated microbe capture accelerates MZ colonization and immune dysfunction by triggering T cell death, T cell-dependent chromatin release and the synergistic induction of G-CSF by histones and fungi. Histones and G-CSF promote the prevalence of immature Ly6G^low^ neutrophils with defective oxidative burst, by selectively shortening the lifespan of mature Ly6G^high^ neutrophils. Consistently, T cell deficiency, or blocking SIGNR1, G-CSF or histones delayed neutrophil dysfunction. Furthermore, histones and G-CSF in the plasma of sepsis patients, shortened neutrophil lifespan and correlated with neutrophil mortality markers associated with a poor prognosis. Hence, the compromise of internal antimicrobial barrier sites drives neutrophil dysfunction by selectively modulating neutrophil lifespan via pathogenic T cell death, extracellular histones, and G-CSF.

## Introduction

Severe sepsis is triggered by systemic infection with micro-organisms that results in aberrant immune responses. It is estimated that in the year 2017 there were 48.9 million sepsis cases, of which 11 million died worldwide, accounting for approximately 20% of all global deaths (*1*). Although fungal-induced sepsis accounts for 5% of microbial sepsis, it is by far the most lethal form with mortality rates exceeding 45% and overall fungal infections cause an estimated 1.6 million deaths world-wide (*2, 3*). The majority of these infections are caused by invasive candidiasis by *Candida albicans*, but the emergence of other pathogenic *Candida* species that are resistant to azoles is of particular concern.

Septic shock is characterized by hyper-inflammation, hypotension, coagulopathy and vascular damage that drive organ failure (*4*). How and where systemic cytokines are induced during sepsis is not well defined. Several damage-associated molecular pattern (DAMP) molecules such as cell-free chromatin, high mobility group protein 1 (HMGB1) and S100 proteins have also been implicated in hyper-inflammation and sepsis pathology (*5–7*). Extracellular histones are cytotoxic and pro-inflammatory (*5, 6, 8, 9*), but their cellular origin and mechanisms regulating their release are unknown.

Immune dysfunction is also prominent during sepsis and is characterised by loss of cytokine responses, T cell deficiency, delayed neutrophil apoptosis and the prominence of immature neutrophils (*10–13*). Neutrophils from patients with septic shock display defective antimicrobial function exemplified by lower oxidative burst capacity (*13–15*). These extreme alterations in neutrophil populations persists well after septic shock and the ensuing immune deficiency negatively impacts survival following an episode of septic shock. Moreover, sepsis patients carry low T cell numbers in their spleens and prominent signs of T cell apoptosis that has been linked to Fas ligand or PD-1/PD-L1 signalling or super-antigen-mediated exhaustion (*16–20*). Similarly delayed apoptosis occurs in mature neutrophils via the triggering receptor expressed on myeloid cells L4 (TREML4) receptor but the signals that regulate the specific loss of mature cells as opposed to immature neutrophils remain unknown (*21*). The signals that drive neutrophil dysfunction and the links with hyperinflammation and immune dysfunction in different immune cell types are poorly understood.

In addition to causing immune deficiency, immune dysfunction may also promote pathology. Neutrophils are critical for controlling invading microbes but are also implicated in tissue destruction and vascular pathology during sepsis (*22*) and SARS-CoV-2 infection pathology (*23*). One antimicrobial protein that is specifically expressed in neutrophils is myeloperoxidase (MPO), a granule enzyme that consumes hydrogen peroxide produced by the NADPH oxidase to generate hypochlorite and other halide oxidants (*24*). Patients with complete MPO deficiencies are prone to recurrent mucosal fungal infections, but rare episodes of systemic fungal infection episodes have also been reported in these patients (*25–27*). MPO deficient mice are more susceptible to pulmonary infection with *C. albicans* and have increased fungal loads upon systemic challenge(*28, 29*). MPO is also required for the release of neutrophils extracellular traps (NETs). NETs control fungi but also promote vascular pathology during sepsis, suggesting that MPO could play beneficial or pathogenic roles during systemic challenge (*27, 29–31*).

Here we show that MPO protects against immune dysfunction by limiting a pathogenic programme that links T cell and neutrophil dysfunction. MPO is critical for controlling infection in the spleen, but over time fungal colonization triggers T cell death that promotes neutrophil dysfunction via the release of cell-free chromatin. Our data demonstrate a pathway that links microbial control to immune dysfunction in the innate and adaptive immune compartments via the critical roles of cell death and inflammatory cytokines.

## Results

### Myeloperoxidase regulates the onset of sepsis

To investigate whether MPO plays a protective or pathogenic role during systemic infection, we performed survival experiments in WT and MPO-deficient mice in a model of systemic candidiasis. WT mice infected with 1×10^5^ WT *C. albicans* yeast succumbed 7-15 days post-infection, whereas MPO-deficient animals developed severe symptoms within 12 hrs post-infection exhibiting a substantial decrease in body temperature (**Fig. 1A and S1A**). In contrast, pulmonary challenge of MPO-deficient animals with a higher fungal load took more than 7 days to lead to serious infection (*29*). MPO-deficient animals did not develop symptoms upon systemic challenge with the yeast-locked Δhgc1 mutant *C. albicans* strain. Moreover, MPO-deficient bone marrow (BM) neutrophils failed to control hyphal germination and growth *in vitro* (**Fig. 1B**). In contrast to human peripheral blood neutrophils, these murine BM cells failed to release NETs as they died with condensed nuclei, indicating that hypochlorous acid but not hydrogen peroxide directly suppresses hyphal growth in the absence of NETosis (**Fig. S1B)**. These experiments suggested that MPO is critical for averting the rapid onset of sepsis.

**Figure 1.**
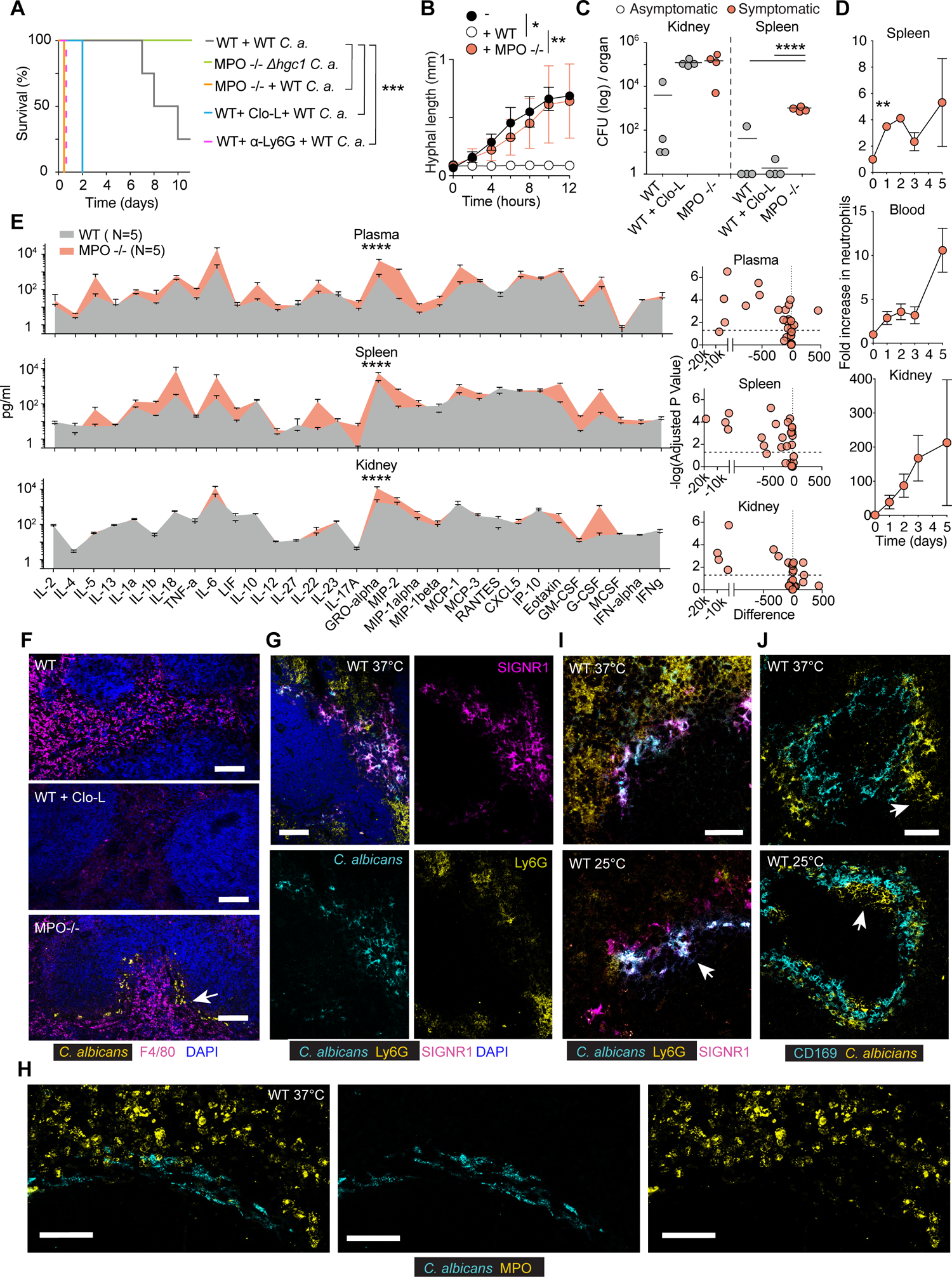
MPO control of fungal colonization in the spleen prevents the rapid onset sepsis. A. Survival curves (left panel) and body temperature (right panel) of WT mice (grey), mice pre-treated with Clo-L (blue), or MPO-deficient mice infected intravenously with 1×10^5^ WT *C. albicans* (orange) or MPO-deficient mice infected with 1×10^5^ *Δhgc1* mutant *C. albicans* (green). 4 experimental replicates with at least 5 mice per group. B. Hyphal growth curves compiled from time-lapse microscopy experiments of WT *C. albicans* (MOI of 0.1) alone (black circles) or in the presence of bone marrow-derived neutrophils from WT (white circles) or MPO-deficient mice (orange circles). Data are represented as mean ± SD. Statistical analysis by Mann-Witney test at 12 hours. C. Colony forming units from homogenized spleens and kidneys isolated 12 hrs post-infection from either WT mice, mice pre-treated with Clo-L, or MPO-deficient mice infected intravenously with 1×10^5^ WT *C. albicans*. Symptomatic mice (temperature lower than 32°C, orange circles) and asymptomatic mice (white open circles). D. Fold increase in Ly6G^+^ neutrophils in the blood, spleens and kidneys of WT mice infected intravenously with 1×10^5^ WT *C. albicans*, before infection and up to 5 days post-infection. E. Cytokine and chemokine protein levels in the plasma, spleen and kidney of WT or MPO-deficient mice infected intravenously with 1×10^5^ WT *C. albicans*, 12 hrs post-infection, assessed by Luminex-based multiplex immunoassay. Statistical analysis Multiple t tests and two-way Anova. Data are represented as mean ± SD. F. Immunofluorescence micrographs from spleens of WT mice, mice pre-treated with Clo-L or MPO-deficient mice infected intravenously with 1×10^5^ WT *C. albicans*, sacrificed 12 hrs post-infection and stained for F4/80, *C. albicans* and DAPI. Arrow points to fungi. Scale bars: 100 μm, data from 2 independent experiments and from at least 4 mice analysed individually. G-J. Immunofluorescence micrographs from spleens of WT mice infected intravenously with 1×10^5^ WT *C. albicans*, 4 - 7 days post-infection with either a body temperature of 37°C or 25°C, stained for (G) *C. albicans*, Ly6G, SIGNR1, (H) *C. albicans* and MPO, (I) *C. albicans*, Ly6G and SIGNR1 or (J) *C. albicans* and CD169. Arrows point to fungi. Scale bars: (G, J, H) 50 μm, (I) 20 μm. data from 2 independent experiments and from with 4 mice analysed individually. Statistical analysis by unpaired Mann-Whitney t-test for single comparisons, Log-rank (Mantel-Cox) test for survival and two-way Anova where indicated (* p<0.05, ** p<0.01, *** p<0.001, **** p<0.0001).

Given that low MPO expression has been reported in macrophages, we also depleted neutrophils with an anti-Ly6G antibody or macrophages with clodronate liposomes (Clo-L)(*32*). Neutrophil depletion resulted in acute symptoms arising within 12 hrs, similarly to MPO-deficient animals. By contrast, Clo-L-treated mice developed symptoms only 48 hrs post-infection (**Fig. 1A**). Consistently, MPO-deficient mice exhibited elevated circulating IL-1β concentrations (**Fig. S1C**) and elevated circulating concentrations of cell-free chromatin **(Fig. S1D and S1E)**. IL-1β and cell-free histones were undetectable in infected WT mice at this early timepoint. The presence of circulating chromatin in MPO-deficient mice confirmed that NETs are not a major source of circulating histones.

To investigate which organs were afflicted by MPO deficiency we examined the fungal load at 12 hrs post-infection. Systemic candidiasis in mice drives sustained high fungal loads in the kidney. By day 1 post-infection, WT mice carried a low kidney fungal load, whereas MPO-deficient and Clo-L-treated mice exhibited comparably elevated fungal load (**Fig. 1C**). However, Clo-L treated animals did not exhibit symptoms of disease as opposed to MPO-deficient animals exhibited significantly higher plasma IL-1β levels and symptoms of severe systemic inflammation such as a hunched position, a scruffy coat, severe hypothermia and lack of responsiveness to stimulation (**Fig. S1A and S1C**). We noted however, that MPO-deficient mice exhibited a significant fungal load in the spleen, suggesting that invasion of this organ may be relevant in the onset of acute symptoms. In symptomatic WT animals, the relative numbers of neutrophils in the spleen peaked rapidly and the total number of cells exceeded the number of neutrophils found in the kidneys (**Fig. 1D and S1F**).

To understand how splenic colonization related to hyperinflammation we analysed pro-inflammatory mediators in the plasma, spleen and kidneys of naïve and infected WT and MPO-deficient animals using luminex-based cytokine arrays. The spleen and blood of MPO-deficient mice exhibited a similar pattern of increase in cytokines and chemokines compared to WT mice (**Fig. 1E**). By contrast, kidney cytokine patterns between infected WT and MPO-deficient mice appeared similar. Together, these findings suggested that neutrophils control splenic colonization and suppress spleen-derived inflammation in an MPO-dependent manner.

Next, we investigated fungal colonization by immunofluorescence microscopy. The kidneys of MPO-deficient and Clo-L-treated mice were colonized throughout the organs (**Fig. S1G**) whereas *C. albicans* colonized predominately the spleen marginal zone (MZ) in MPO-deficient animals (**Fig. 1F**). The spleen contains several different macrophage populations. MZ macrophages are comprised of the outer layer that expresses the receptors SIGN-related 1 (SIGNR1) and MARCO and the inner layer expresses CD169 (**Fig. S1H**). Clo-L administration had little effect on splenic macrophages within the first 24 hrs, but 48 hrs post-treatment eliminated a substantial number of F4/80^+^ red pulp cells, depleted SIGNR^+^ macrophages completely and had a partial effect on CD169^+^ cells leaving a substantial number of CD169^+^ MZ macrophages were still present after Clo-L treatment (**Fig. S1I**). WT and Clo-L-treated mice did not bear visible microbial colonies in the spleen 24 hrs post-infection which correlated with the lack of symptoms in these mice (**Fig. 1F**).

To determine the precise localization of fungi in the spleen, we stained the spleens of infected WT mice collected from a period ranging from 4-7 days post-infection, which contained asymptomatic mice and mice that developed sepsis symptoms. Fungi colonized the spleen MZ of WT mice and colocalized with SIGNR1^+^ macrophages in asymptomatic mice with physiological body temperature 37°C (**Fig. 1G**). *C. albicans* and Ly6G+/MPO^+^ neutrophils did not co-localize in the spleen (**Fig. 1G and 1H**). In WT mice that had developed severe symptoms and exhibited a low body temperature, *C. albicans* had infiltrated the inner CD169^+^ macrophage layer, suggesting that spreading of the microbes through this barrier may play a role in pathology (**Fig. 1I and 1J**). Taken together, these data indicated that neutrophils were critical to control fungi that were captured by SIGNR1^+^ macrophages via MPO-derived ROS. The fact that MPO delayed colonization in areas that did not contain neutrophils suggested that neutrophil-derived ROS controlled hyphal growth in a non-cell-autonomous, transcellular manner.

### SIGNR1-mediated capture promotes fungal colonization and cytokine production at the spleen MZ

Next, we explored whether the capture of fungi promotes sepsis pathology by targeting the SIGNR1 receptor. The C-type lectin receptor SIGNR1 is predominately expressed in the spleen and lymph nodes and binds *C. albicans* (*33*). SIGNR1^+^ cells could not be detected in the infected kidneys indicating that the antibody would not directly impact immune responses in this organ (**Fig. S2A**). Moreover, we confirmed that this antibody did not deplete SIGNR1^+^ macrophages because we could still detect them in naïve mice with antibodies against MARCO (**Fig. S2B**). SIGNR1 staining was not detectable in the spleens of mice treated with anti-SIGNR1, indicating the effective blockade of the receptor (**Fig. S2C**). For these experiments, we raised the infection dose to 5×10^5^ yeast particles per animal in order to facilitate the detection of any survival benefit against untreated WT mice that are more resistant to infection than MPO knockout animals. With the higher fungal dose, WT mice treated with a control IgG antibody developed severe symptoms 3 days post-infection, whereas SIGNR1 blockade delayed the onset of symptoms and extended survival by 3-fold (**Fig. 2A and S2D**). SIGNR1 blockade reduced the fungal load in the spleen as well as the kidneys, despite the receptor not being expressed in this organ (**Fig. 2B and S2E**). Moreover, SIGNR1 blockade reduced NET deposition in the kidneys (**Fig. S2F**) and lowered cytokines such as IL-6 and G-CSF and chemokines in the spleen, blood and kidney 3 days post-infection (**Fig. 2C and S2G**). To determine which splenocytes produced cytokines we detected cytokine mRNA transcripts by in fluorescence situ hybridization (RNAscope) staining in splenic sections. IL-6 and G-CSF were upregulated in CD169^+^ macrophages in infected IgG-treated mice and were reduced in anti-SIGNR1 treated mice (**Fig. 2D**). Hence, SIGNR1 accelerates the onset of sepsis pathology and exerts its effects systemically in organs lacking cells that express the receptor.

**Figure 2.**
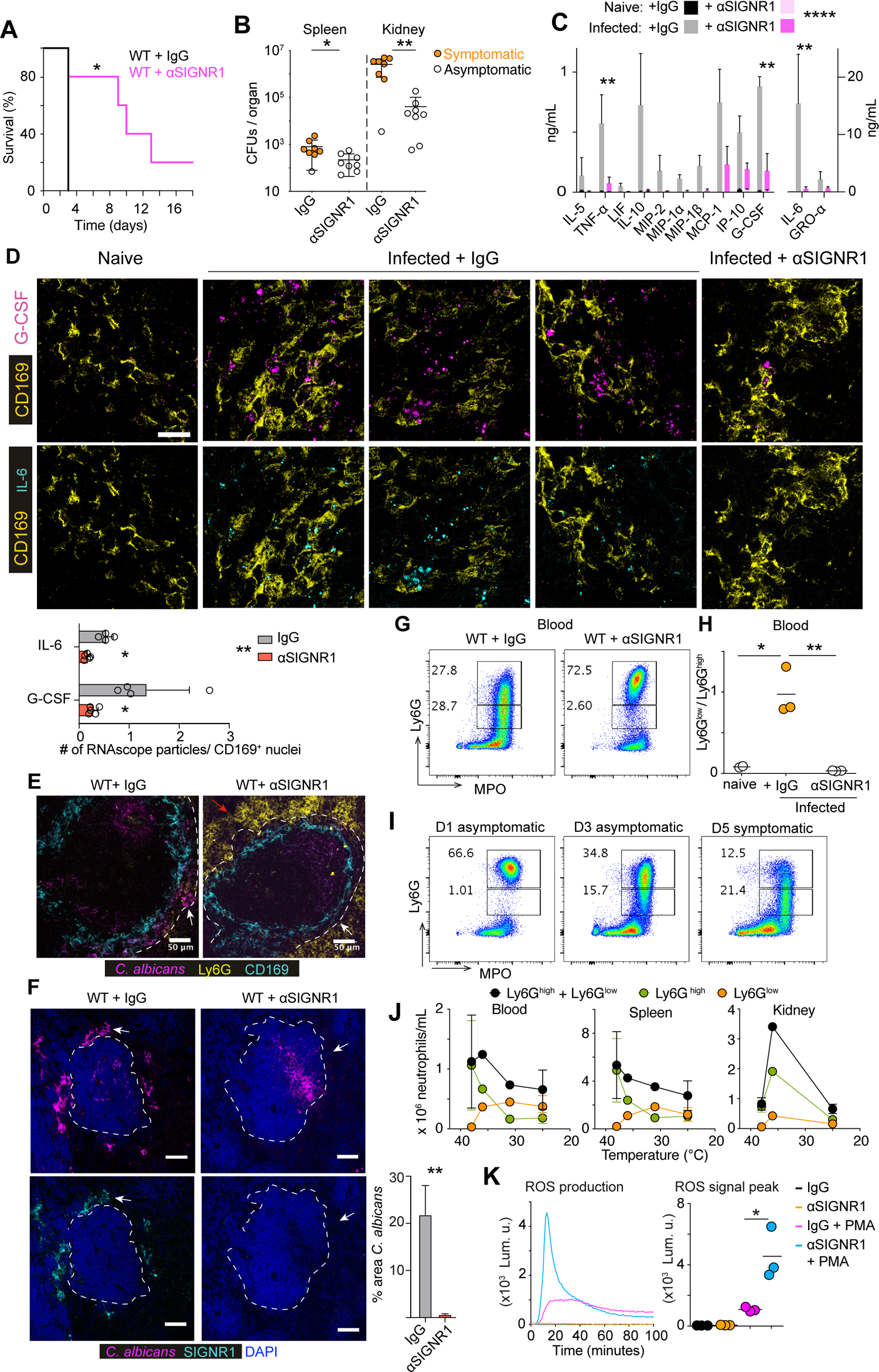
SINGR1 regulates fungal colonization and cytokine production and altered ROS-deficient circulating neutrophils. A-J. WT mice pre-treated with either control antibody or an anti-SIGNR1 blocking antibody and infected intravenously with 5×10^5^ WT *C. albicans*: A. Survival curves. B. Colony forming units 72 hrs post-infection. Symptomatic mice are shown in orange whereas asymptomatic mice in white open circles. C. Selected cytokines in plasma 72 hrs post-infection detected by Luminex-based multiplex immunoassay. D. Representative RNA scope imaging of G-CSF and IL-6 expression in the spleens 72 hrs post-infection co-stained with anti-CD169 antibodies, imaged by immunofluorescence confocal microscopy. Scale bars: 20 μm, Representative results from 5 mice per group. The average number of RNA scope particles / CD169 positive nuclei (DAPI) in images from 4 mice per group is shown below. Statistical analysis by unpaired Mann-Whitney t-test for pairwise comparison and two-way Anova for the experiment. E and F. Immunofluorescence confocal micrographs of spleens from mice treated with control or anti-SIGNR1 antibody and infected with 5×10^5^ WT *C. albicans*, 3 days post-infection and stained for (E) *C. albicans*, Ly6G, CD169, or (F) C. albicans, SIGNR1 and DAPI. Arrows point to the MZ. Scale bars: 50 μm. The panel on the bottom right corner shows the fraction of MZ area occupied by *C. albicans*. G. Flow cytometric analysis of neutrophils isolated from peripheral blood of mice treated with IgG control or anti-SIGNR1 antibodies and infected with 5×10^5^ WT *C. albicans*, 3 days post-infection stained for intracellular MPO and Ly6G expression. The gate on top indicates the percentage of Ly6G^high^ neutrophils and the gate below the Ly6G^low^ neutrophils. The right panel depicts Ly6G^low^/Ly6G^high^ neutrophils ratios per animal. Gates were set on Live CD3^-^ CD19^-^ CD11b^+^ cells. H. Quantification of Ly6G^low^/Ly6G^high^ ratios from (G) in the blood of individual mice. I. Flow cytometric analysis of neutrophils isolated from peripheral blood of WT mice infected with 5×10^5^ WT *C. albicans,* 1, 3 and 5 days post-infection. Cells were stained for intracellular MPO and surface Ly6G expression. N=3-5 mice per group in 2 independent experiments. J. Plots depicting total neutrophil numbers in the blood, spleen and kidney of infected WT mice against their body temperature. K. ROS production in the absence or presence of PMA, by neutrophils isolated 3 days post-infection from the spleens of mice that were pre-treated with either IgG control or anti-SIGNR1 antibody and infected with 5×10^5^ WT *C. albicans*. A representative curve from one mouse per group depicting ROS detected by luminol bioluminescence over time post-stimulation (left) and a cumulative plot of the ROS peak level for the neutrophils from each animal (right). Statistical analysis by unpaired Mann-Whitney t-test for single comparison and Log-rank (Mantel-Cox) test for survival analysis (* p<0.05, ** p<0.01, *** p<0.001, **** p<0.0001).

### SIGNR1 regulates neutrophil populations

Next, we investigated the impact of SIGNR1 blockade on fungal capture in the spleen. Anti-SIGNR1 treatment prevented the sequestration of *C. albicans* in the spleen MZ but microbes could be detected in other splenic areas, such as the white pulp. (**Fig. 2E and 2F**). In addition, we also noticed that Ly6G staining disappeared in infected IgG-treated mice but was present in anti-SIGNR1-treated animals (**Fig. 2E**). The prominent MPO staining in the infected spleens of untreated WT mice (**Fig. 1I**) suggested that neutrophils were still present in large numbers but may have downregulated Ly6G expression. To further test this hypothesis, we analysed neutrophil populations in the blood, spleens and kidneys by flow cytometry, tracking CD11b, MPO and Ly6G. The administration of the anti-SIGNR1 antibody did not alter the total neutrophil numbers in the blood on the day of infection nor did it affect emergency granulopoiesis 24 hrs post-infection (**Fig. S3A**). However, 3 days post-infection, control animals contained a large fraction of neutrophils that had high MPO but low Ly6G expression (**Fig. 2G and 2H**). In contrast, mice treated with anti-SIGNR1 maintained the Ly6G^high^ population in the blood, spleen and kidneys (**Fig. 2G, 2H and S3B**). The shift towards an elevated Ly6G^low^/ Ly6G^high^ neutrophil ratio in the blood, spleen and kidney was gradual and preceded the onset of sepsis symptoms (**Fig. 2I and S3C and S3D**). We also plotted the change in neutrophil populations against the corresponding body temperature in an experiment where WT mice became gradually sick and were sacrificed at an intermediate timepoint that contained a mixed group of symptomatic and asymptomatic mice. These plots showed that although the total neutrophil population decreased by approximately 30%, there was a near 10-fold decrease in Ly6G^high^ neutrophils and a trend for an increase in Ly6G^low^ cells as body temperature decreased (**Fig. 2J**).

Next, we explored whether the shift in neutrophil populations influenced the antimicrobial capacity of neutrophils. We evaluated superoxide production in neutrophils isolated from the spleens of infected control and anti-SIGNR1-treated mice 3 days post-infection. Neutrophils from anti-SIGNR1-treated mice produced a potent ROS burst response to phorbol-myristate-acetate (PMA), whereas neutrophils from control-treated mice exhibited a defective ROS burst (**Fig. 2K**). Interestingly, SIGNR1 blockade failed to improve the survival of infected MPO-deficient mice even at 100-fold lower infection doses confirming that the beneficial impact of SIGNR1 blockade is mediated by regulating neutrophil effector function (**Fig. S3E**). These data indicated that SIGNR1-mediated fungal capture controls splenic colonization and neutrophil dysfunction.

### SIGNR1 promotes T cell death and chromatin release

The presence of cytokines in MZ macrophages raised the question of whether they are undergoing cell death. Hence, we stained infected spleens with terminal deoxynucleotidyl transferase dUTP nick end labelling (TUNEL). We detected no substantial TUNEL signal in MZ macrophages but there was extensive death of cells in the white pulp of infected control-treated mice (**Fig. 3A**). Notably, cell death was absent in SIGNR1-blocked mice and staining with T cell markers demonstrated that the dying cells were predominately T cells that were positive for CD4, CD3 and TCRβ (**Fig. 3A, 3B and S4A**). Interestingly, a substantial number of TUNEL^+^ T cells could be found outside germinal centers (**Fig. 3A, 3C and S4A**). We also confirmed the loss of T cells in the spleen by flow cytometry which demonstrated that T cell death impacted T cells non-specifically and afflicted all Vβ-chain populations (**Fig. 3D**). Staining these splenic samples with antibodies against the activated form of caspase-3 and capsase-8 confirmed that T cells were dying via apoptosis (**Fig. 3E**). We also detected a SIGNR1-dependent increase in T cell death in the inguinal lymph nodes and the thymus although the latter was not statistically significant (**Fig. 3F, 3G and S4B**). Interestingly, *C. albicans* was absent from lymph nodes and the thymus, indicating that T cell death was driven by systemic mediators (**Fig. S4C**). Cell death also occurred in the kidneys and was significantly reduced in SIGNR1-blocked mice. However, dying kidney cells were not T cells as TUNEL positive cells were not positive for CD3 (**Fig. S4D**). Given the extent of cell death, we examined whether SIGNR1 blockade affected circulating cell-free chromatin. Consistently, DNA and histones were prominent in the plasma of infected IgG-treated mice but were absent in samples from SIGNR1-treated mice (**Fig. 3H and 3I**). Collectively, these data demonstrate that SIGNR1 regulates T cell death and the release of extracellular chromatin in the circulation.

**Figure 3.**
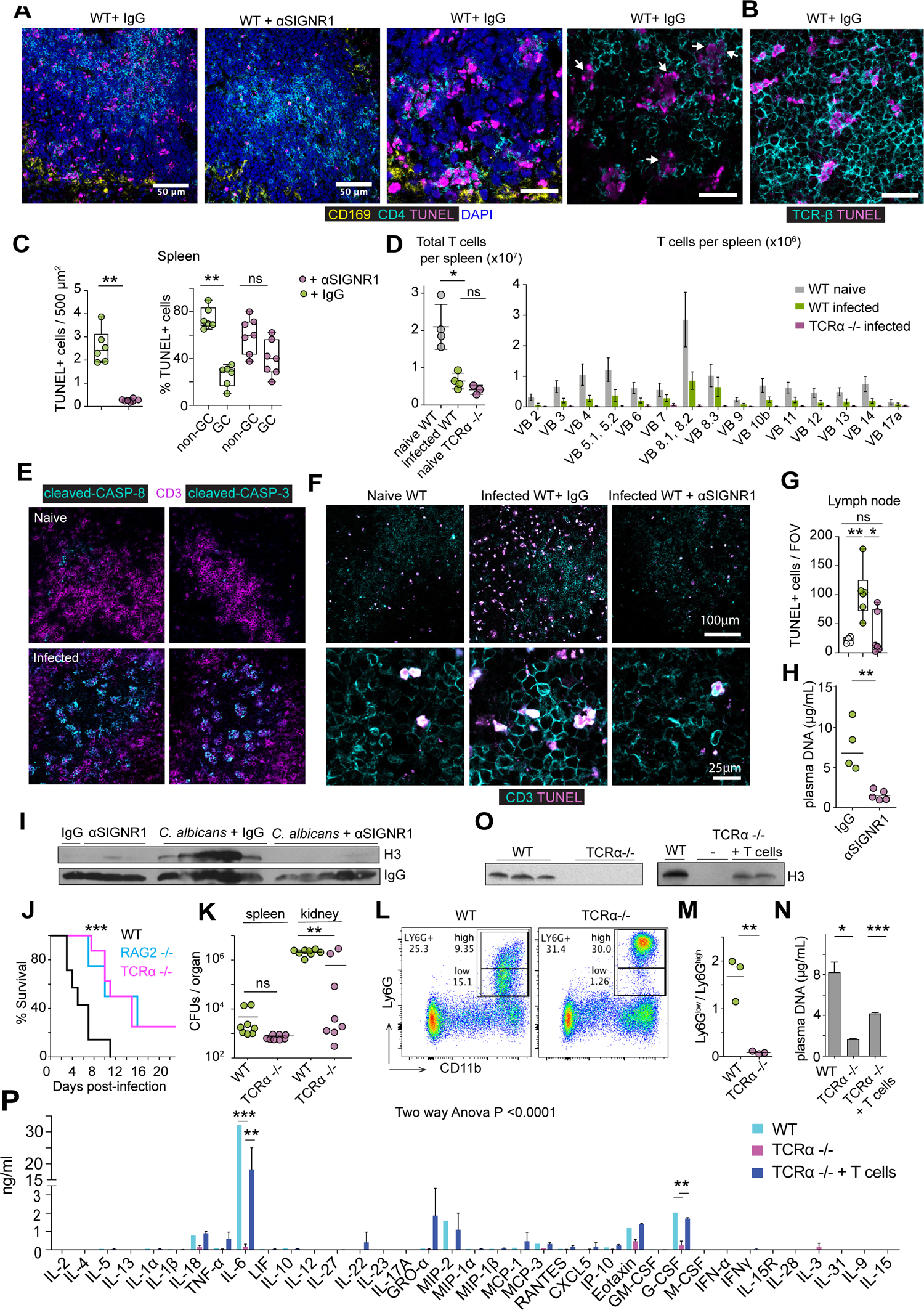
SINGR1 regulates T cell death, circulating chromatin release and neutrophil alterations. A. Immunofluorescence micrographs from spleens of WT mice infected with 5×10^5^ WT *C. albicans* 3 days post-infection that were pre-treated with either control or anti-SIGNR1 antibody and stained for CD169, TUNEL, CD4 and DAPI. Scale bars: 50 μm (left panels) and 20 μm (right panels). B. Staining of spleens from symptomatic WT infected animals with TUNEL and anti-TCR-β antibodies. Scale bar: 20 μm. C. Quantification of TUNEL^+^ cells in the white pulp of the spleen per 500 μm^2^ (left panel), and distribution of TUNEL^+^ positive cells within or outside germinal centers (GC). D. Flow cytometric analysis of the numbers of total T cells (left) and TCR Vβ subset T cells (right) in the spleens of naïve WT, naive TCRα-deficient mice (negative control) and WT mice infected with 5×105 WT C. albicans, 2 days post-infection. Gates were set on Live CD19-CD3+ cells. E. Immunofluorescent micrographs from spleens of naïve WT mice or infected with 5×10^5^ WT *C. albicans* 3 days post-infection stained for CD3 and antibodies specific for the cleaved activated form of either caspase-3 or caspase-8. F. Immunofluorescent micrographs of inguinal lymph nodes from mice in (A) stained for TUNEL and CD3 at lower magnification (upper row) scale bars 100 μm, and higher magnification scale bar 25 μm (bottom row). G. Quantification of TUNEL+ cells per field of view (FOV) in inguinal lymph nodes and thymus. Three FOVs per mouse and 3 mice per group were analysed. H and I. Cell-free DNA (H) and histone H3 (I) and in the plasma of WT mice pre-treated with either control or anti-SIGNR1 antibody and infected with 5×10^5^ WT *C. albicans*, 3 days post-infection. J. Survival curves of WT, TCRα or RAG2 deficient mice infected with 5×10^5^ *C. albicans*. N=5 mice per group K. Colony forming units in the spleens and kidneys of WT and TCRα-deficient mice infected with 5×10^5^ *C. albicans*. Representative of two experimental replicates. L. Representative flow cytometric analysis of Ly6G^low^ and Ly6G^high^ neutrophils in the spleen of WT and TCRα-deficient mice infected with 5×10^5^ WT *C. albicans* and assessed as Live CD3^-^ CD19^-^ CD11b^+^ Ly6G^high^/Ly6G^low^ 72 hrs post-infection. M. Ly6G^high^/Ly6G^low^ neutrophil ratios in individual mice from (L). N-P. Cell-free DNA (N), western immunoblotting of histone H3 (O), and plasma cytokines and chemokines measured by multiplex immunoassay (P) in the plasma of WT and TCRα-deficient mice alone or after adoptive transfer of T cells and infected with 5×10^5^ WT *C. albicans*. Samples were harvested 72 hrs post-infection. N=3-5 mice group, results from 2-3 independent experiments. Statistical analysis by unpaired Mann-Whitney t-test for single comparisons and the Log-rank (Mantel-Cox) test for survival analysis (* p<0.05, ** p<0.01, *** p<0.001, **** p<0.0001).

### T cells regulate circulating chromatin release and neutrophil populations

To evaluate whether T cell death was pathogenic we examined the survival of RAG2-deficient mice that lack T cells and B cells and TCRα-deficient mice that lack αβ-T cells. Both knockout strains exhibited improved survival (**Fig. 3J**) and carried a lower fungal load in their kidneys 3 days post-infection (**Fig. 3K**). Moreover, infected TCRα-deficient mice maintained Ly6G^high^ neutrophils in the periphery indicating that T cells are involved in neutrophil dysfunction (**Fig. 3L and 3M**). Moreover, TUNEL staining was absent from the spleens of infected TCRα-deficient mice and adoptive transfer of T cells restored cell death in the spleen white pulp (**Fig. S4E**). Staining with a FLICA poly-caspase activity assay further confirmed the elevated incidence of cell death in T cells and was absent in the spleen of T cell deficient mice (**Fig. S4F**). Likewise, circulating chromatin was absent in the plasma of TCRα knockout animals and could be restored by adoptive transfer of T cells (**Fig. 3N and 3O**). Furthermore, T cell deficiency led to a reduction in plasma cytokines, including IL-6 and G-CSF (**Fig. 3P**). Together, these observations indicated that dying T cells regulated the release of circulating chromatin, cytokine induction and alterations in neutrophil populations.

### Cell death accelerates alterations in neutrophil populations

Next, we investigated whether cell death plays a role in cytokine induction and neutrophil dysfunction. Macrophage depletion experiments using Clo-L administration induced very high concentrations of circulating chromatin in the absence of infection but did not cause pathology (**Fig. 4A**). The spleen is an organ rich in macrophages, therefore we hypothesized that the phenotype we observed in infected Clo-L-treated mice was not only caused by a reduction in macrophages but could also be accelerated by pre-existing chromatin release. Consistent with this hypothesis, Clo-L treatment accelerated the emergence of Ly6G^low^ neutrophils and the reduction in total splenic neutrophil numbers upon infection (**Fig. 4B and S5A**). Therefore, cell death and circulating histones alone were not sufficient to induce pathology in the absence of infection but accelerated cytokine induction and neutrophil alterations upon infection.

**Figure 4.**
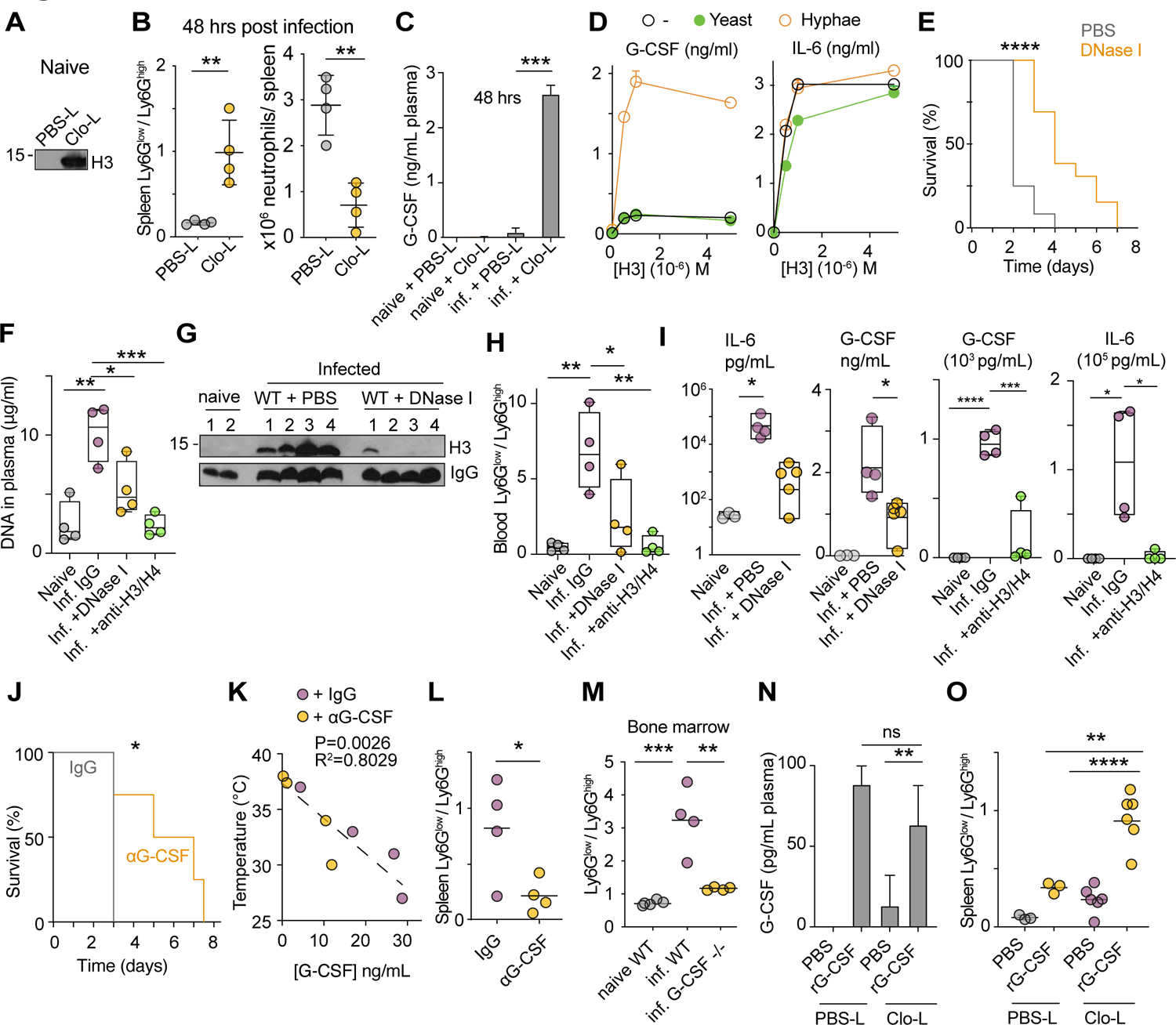
Circulating chromatin and G-CSF regulate neutrophil dysfunction. A. Western immunoblotting for histone H3 in the plasma of naïve WT mice 24 hrs after intravenous injection of PBL-liposomes (PBS-L) or clodronate-liposomes (Clo-L). B. Ly6G^low^/Ly6G^high^ ratios (left panel) and total neutrophils (right panel) in the spleens of naïve WT mice pre-treated with PBS-L or Clo-L, or pre-treated mice infected with 1×10^5^ WT *C. albicans,* harvested 48 hrs post-infection. C. Plasma G-CSF concentrations assessed by simplex immunoassay in naïve mice and WT mice pre-treated PBS-L or Clo-L, either naïve or infected with 1×10^5^ WT *C. albicans,* 48 hrs post-infection. D. G-CSF (left) and IL-6 (right) protein production in WT bone marrow-derived macrophages alone (open black) or in the presence of heat-inactivated yeast (solid green) or hyphae (open orange) and in combination with increasing concentrations of recombinant histone H3. E. Survival of WT mice pre-treated with vehicle (PBS, grey) or DNase I (orange) and infected with 5×10^5^ WT *C. albicans*. (F-G) Cell free DNA levels (F) and Western immunoblot of histone H3 (G) from the plasma of naïve WT mice or infected with 5×10^5^ WT *C. albicans* and pre-treated with either IgG control antibody, anti-H3 and anti-H4 antibodies or DNase I, analysed 72 hrs post-infection. H. IL-6 and G-CSF in the plasma of mice in (F) measured by multiplex immunoassay. I. Ly6G^low^/Ly6G^high^ ratios in splenic neutrophils from mice in (E). J. Survival of WT mice infected with 5×10^5^ WT *C. albicans* and treated daily with control antibody or anti-G-CSF antibody starting 48 hrs post-infection. K. Correlation between plasma G-CSF concentrations and body temperature at 3 days post-infection in WT mice infected with 5×10^5^ *C. albicans* and treated with control antibody (open circles) or anti-G-CSF antibody (orange circles), started 2 days post-infection. L. Ly6G^low^/Ly6G^high^ ratios in splenic neutrophils from mice in (K) 3 days post-infection. M. Neutrophils analysed from the bone marrow of WT and G-CSF deficient FVB/NJ mice, either naïve or infected with 1×10^4^ or 1×10^3^ WT *C. albicans* respectively and measured on the day severe symptoms appeared based on the decrease in body temperature below 32 °C. N. Plasma G-CSF concentrations in naïve mice treated with PBS-L or Clo-L, subsequently injected with PBS or recombinant G-CSF (rG-CSF) after 24 hrs and assessed 48 hrs later. O. Ly6G^low^/Ly6G^high^ ratios in splenic neutrophils from mice in (N). N=3-6 mice per group. Statistical analysis by unpaired Mann-Whitney t-test for single comparison and Log-rank (Mantel-Cox) test for survival analysis (* p<0.05, ** p<0.01, *** p<0.001, **** p<0.0001).

### Circulating chromatin and fungi induce G-CSF

Since there was a notable increase in G-CSF that correlated with the emergence of Ly6G^low^ neutrophils, we examined whether G-CSF was differentially induced upon infection in Clo-L treated mice. While Clo-L alone in the absence of infection had little effect on the plasma concentrations of G-CSF, Clo-L pre-treatment amplified circulating G-CSF concentrations upon fungal challenge (**Fig. 4C**). These data suggested that circulating histones and *C. albicans* could synergize *in vivo* to induce G-CSF. To test this hypothesis, we incubated bone marrow-derived macrophages with recombinant histone H3 alone or in combination with heat-inactivated yeast or hyphae. Histones or hyphae alone were weak G-CSF inducers, but the two signals acted synergistically to induce the cytokine (**Fig. 4D**). This synergy depended on the hyphal form, as yeast were unable to induce G-CSF in the presence of histone H3. By contrast, histone H3 was sufficient to induce IL-6 and hyphae had no effect on the induction of this cytokine suggesting a divergence in the pathways that regulate IL-6 and G-CSF.

### Circulating chromatin and G-CSF are required to alter neutrophil populations

To test the impact of histones on neutrophils we sought an approach that would clear circulating chromatin enzymatically. We previously found that proteases and endonucleases synergize in chromatin clearance in the sputum of patients with cystic fibrosis (*34*). Therefore, we tested whether intraperitoneal administration of DNase I could promote chromatin clearance in the plasma that contains serum proteases (*5*). We also treated mice with histone-blocking antibodies. DNase I administration delayed sepsis in infected mice (**Fig. 4E**) and cleared free circulating DNA and histones (**Fig. 4F and 4G**). Moreover, mice treated with either DNase I or anti-histone antibodies maintained high circulating Ly6G^high^ neutrophil populations (**Fig. 4H and S5B**) and decreased plasma G-CSF and IL-6 concentrations (**Fig. 4I, S5C and S6A**) suggesting that plasma chromatin may be affecting neutrophil populations by regulating cytokines. Histone-blocking was more effective than DNAse I treatment and completely suppressed the increase in the Ly6G^low^/Ly6G^high^ ratio as well as maintained a physiological body temperature (**Fig. S6B**).

The link between cell-death-derived chromatin and G-CSF was intriguing since this cytokine is a key neutrophil regulator that amplifies granulopoiesis and mobilizes neutrophils from the bone marrow. G-CSFR-deficient mice are more susceptible to systemic *C. albicans* challenge despite exhibiting effective emergency granulopoiesis and neutrophil mobilization from the bone marrow (*35*). However, in our experiments G-CSF correlated with neutrophil dysfunction and was upregulated by SIGNR1 and circulating chromatin. Therefore, we hypothesized that prolonged exposure to G-CSF may promote pathology during systemic infection. To test this hypothesis, we neutralized G-CSF by daily administration of antibodies, starting at 24 - 48 hrs post-infection, in order to avoid interfering with any protective functions of G-CSF during the early phases of the infection. Late anti-G-CSF treatment resulted in a significant delay in the onset of severe symptoms (**Fig. 4J**). Elevated G-CSF plasma concentrations correlated with a decrease in body temperature (**Fig. 4K**) and G-CSF neutralization suppressed the emergence of Ly6G^low^ neutrophils (**Fig. 4L and S7A**). We also infected G-CSF-deficient mice with *C. albicans* and examined the effect on neutrophil populations. Due to peripheral neutropenia these mice are more susceptible to *Candida* systemic infection and therefore we infected with the low 1×10^3^ *C. albicans* inoculum which induced severe symptoms 2-3 days post-infection. Given that G-CSF elicits neutrophil egress from the bone marrow we analysed neutrophils in the bone marrow to ensure that any measured differences were not due to lack of neutrophil mobilization. Notably, the Ly6G^low^/Ly6G^high^ neutrophil ratio in infected symptomatic G-CSF-deficient mice remained low (**Fig. 4M**) suggesting that G-CSF is required for the observed alterations in neutrophil populations.

### Cell death and G-CSF are sufficient to alter neutrophil populations in the absence of infection

The administration of G-CSF in the absence of infection is not known to cause large deleterious effects on granulopoiesis. However, G-CSF is required but not sufficient to upregulate minor populations of low-density granulocytes in murine cancer models (*36*). Therefore, we hypothesized that G-CSF may synergize with cell death-derived chromatin in regulating neutrophil dysfunction. Moreover, we sought conditions where we could decouple the emergence of Ly6G^low^ neutrophils from infection. To test this hypothesis, we treated mice with PBS liposomes (PBS-L) or Clo-L and injected recombinant G-CSF after 24 hrs. We assessed splenic neutrophil populations by flow cytometry 48 hrs later. At that time, PBS-L and Clo-L-treated groups that had received rG-CSF had comparable G-CSF concentrations in their blood (**Fig. 4N**). However, while G-CSF injection in PBS-L-treated mice did not alter the neutrophil ratio substantially, the combination of Clo-L and G-CSF induced a Ly6G^low^/Ly6G^high^ neutrophil ratio that was comparable to that observed in infected symptomatic mice (**Fig. 4O and S7B**). Pre-treating mice with recombinant G-CSF 24 hrs prior to infection delayed the onset of sepsis indicating that boosting granulopoiesis prior to infection is beneficial and that G-CSF plays distinct roles in the presence and in the absence of circulating chromatin (**Fig. S7C**). Together these data indicated that in addition to the ability of histones to induce G-CSF in the presence of systemic infection, chromatin also acts downstream with G-CSF to interfere with the antimicrobial fitness of neutrophil populations.

### The transcriptional signature of Ly6G^low^ neutrophils resembles immature populations and is regulated by T cells

To better understand the changes in neutrophil populations during sepsis and the impact of T cell death, we profiled neutrophils populations. Wright-Giemsa staining indicated that FACS-sorted Ly6G^low^ neutrophils from symptomatic mice, comprised a mixed population of mature lobulated and morphologically immature band cells (**Fig. 5A**). To compare these cells at the molecular level, we conducted RNA sequencing analysis of Ly6G^high^ and Ly6G^low^ neutrophils sorted by flow cytometry from the spleens of either naïve or infected WT or TCRα-deficient animals and uninfected WT mice treated with G-CSF and Clo-L. PCA analysis indicated that Ly6G^high^ and Ly6G^low^ populations from infected WT symptomatic mice were distinct from one another but also clustered differently from Ly6G^high^ neutrophils isolated from infected asymptomatic TCRα-deficient animals or naïve WT mice (**Fig. 5B**). Moreover, Ly6G^low^ neutrophils in G-CSF and Clo-L treated mice were distinct from naïve Ly6G^high^ cells but also exhibited variations from Ly6G^low^ neutrophils found in infected symptomatic mice. Next, we compared these populations based on the major gene expression differences between septic Ly6G^low^ and Ly6G^high^ cells across the entire dataset. Ly6G^low^ neutrophils from infected WT mice had a distinct signature in comparison to Ly6G^high^ populations but also when compared to Ly6G^low^ neutrophils in infected asymptomatic TCRα-deficient mice (**Fig. 5C**). Given the asymptomatic state, the TCRα-deficient Ly6G^low^ cells comprised a “healthy” immature population under a state of infection. TCRα-deficient mice exhibited a Ly6G^high^ population that was similar to Ly6G^high^ populations in naïve WT mice. Moreover, WT infected Ly6G^low^ cells were closer to Ly6G^low^ cells from TCRα-deficient mice but also exhibited distinct transcriptional changes that set them apart from immature cells associated with an infected asymptomatic state.

**Figure 5.**
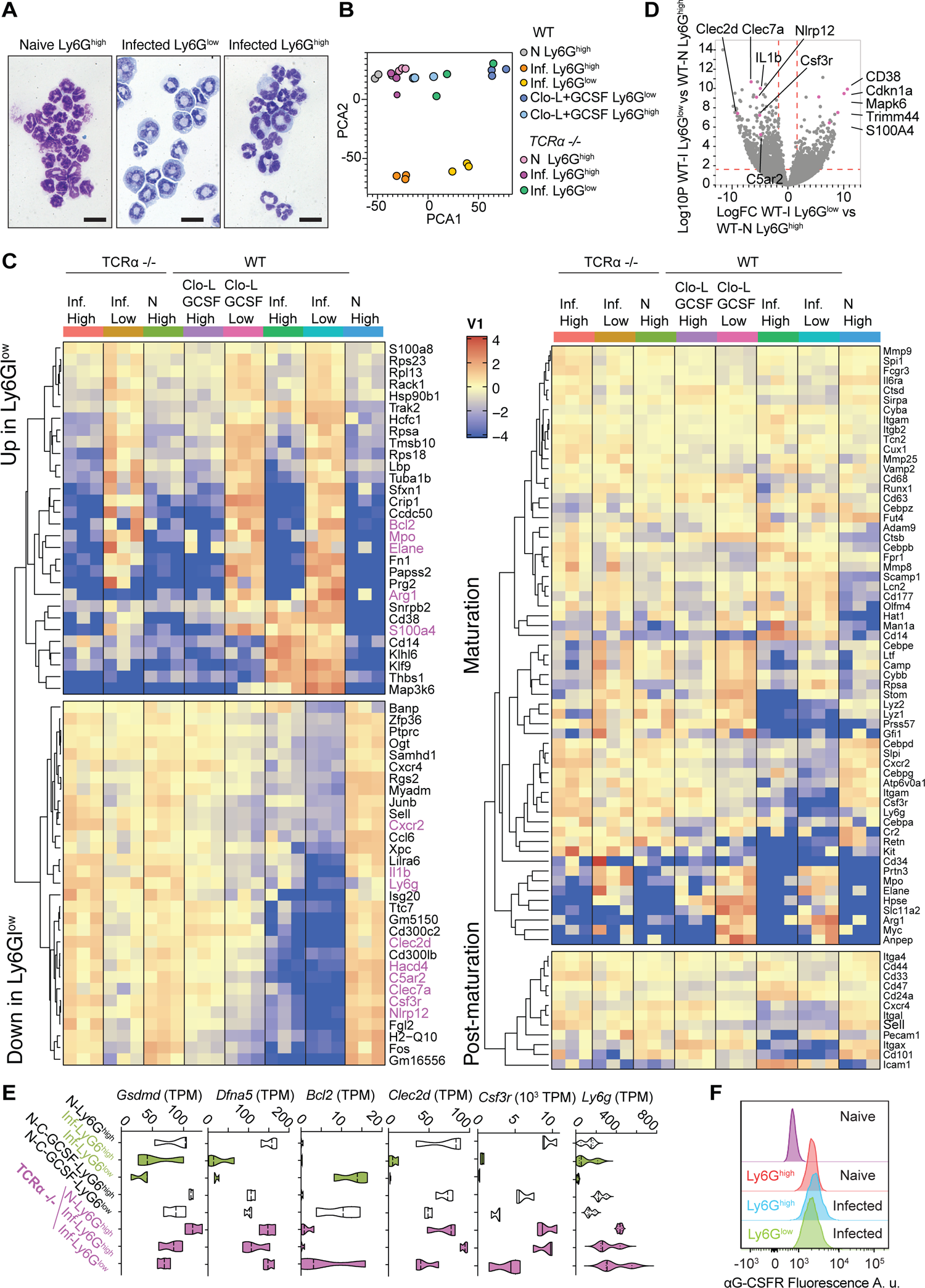
Transcriptomic profiling of neutrophils during infection and sterile challenge. A. Giemsa-Wright staining of Ly6G^low^ and Ly6G^high^ neutrophils from WT naïve mice or infected with 5×10^5^ WT *C. albicans,* 72 hrs post-infection, that were cyto-spun following sorting by flow cytometry. Sorted cells were LiveCD3^-^CD19^-^CD11b^+^Ly6G^high/low^. B. Principal component analysis of transcripts from sorted Ly6G^low^ and Ly6G^high^ neutrophils isolated from WT and TCRα-deficient mice, either naïve or infected with 5×10^5^ WT *C. albicans* (48 hrs post-infection) and naïve WT mice treated with Clo-L and recombinant G-CSF. C. Expression profiling heatmaps for the (left) top genes that are differentially expressed and (right) classical maturation and post-maturation markers in neutrophils from all groups in (B). D. Volcano plot comparing expression profiling of flow cytometric sorted Ly6G^high^ and Ly6G^low^ neutrophils from WT naïve and infected mice respectively, 48 hrs post-infection. E. Transcript reads per million (TPM) of Gsdmd, Dfna5, Bcl2, Clec2d, Csf3r and Ly6g. F. Flow cytometric analysis of G-CSFR protein surface expression in Ly6G^high^ neutrophils from naïve mice and Ly6G^high^ and Ly6G^low^ neutrophils from mice infected with 5×10^5^ WT *C. albicans*, 72 hrs to 96 hrs post-infection. Includes an FMO control containing only secondary antibody. N=4-5 mice per group. Statistical analysis by unpaired Mann-Whitney t-test (* p<0.05, ** p<0.01, *** p<0.001, **** p<0.0001).

There was a cluster of genes with similar trends amongst the uninfected Ly6G^low^ cells isolated from uninfected G-CSF/Clo-L-treated-mice and the Ly6^low^ neutrophils derived from infected symptomatic WT mice supporting the hypothesis that G-CSF and chromatin promote shifts in neutrophil populations during infection. We also compared these cell populations for genes associated with the different stages of neutrophil maturation and found that Ly6G^low^ neutrophils exhibited reduced expression in several maturation and post-maturation genes (**Fig. 5C**). Comparison of infected WT Ly6G^low^ with Ly6G^low^ neutrophils from TCRα-deficient mice also highlighted changes in a number of genes that included CXCR2, CD14 ITGAM, Csf3r, CD34, CR2 and Arg1, which distinguished this population from TCRα-deficient Ly6G^low^ cells. Infected WT Ly6G^low^ neutrophils also lacked transcripts for C-type-lectin-receptor Clec2d and Clec7a, two C-type lectin receptors that recognize histones and fungal pathogens respectively as well as the complement receptor C5ar2, two Nod-like receptors and IL-1β, which could further compromise their antifungal activity (**Fig. 5C and 5D**). Notably, while Ly6g transcripts were downregulated in Ly6G^low^ cells than in Ly6G^high^ neutrophils from infected WT spleens, the transcripts for the receptor were comparably high in both populations isolated from infected TCRα-deficient mice (**Fig. 5E**), although this was not reflected at the protein level. Infected WT Ly6G^low^ cells had additional adaptations that distinguished them from TCRα-deficient Ly6G^low^ neutrophils such as low levels of CXCR2, gasdermin D and E (Gsdmd and Dfna5) and higher Bcl2 transcript levels which may increase cell survival. Moreover, transcripts for the G-CSF receptor Csf3r were downregulated in Ly6G^low^ neutrophils from infected mice or uninfected mice treated with G-CSF and Clo-L. The downregulation of G-CSFR mRNA could result from a negative feedback response to sustained G-CSF signalling in WT infected animals that was absent in TCRα-deficient mice. However, the G-CSFR protein could still be detected on the surface of Ly6G^high^ and Ly6G^low^ neutrophils from infected WT mice at comparable intensity as in Ly6G^high^ cells from naïve WT mice indicating that these cells were still sensitive to G-CSF (**Fig. 5F and S7D**). These data confirmed that peripheral neutrophil populations in symptomatic mice were predominately immature cells that were molecularly distinct from peripheral mature and immature neutrophils found in infected mice lacking T cells, indicating that additional changes occur with disease severity.

### T cell-derived histones eliminate mature Ly6G^high^ neutrophils but not Ly6G^low^ neutrophils in the bone marrow

To explore the underlying mechanism behind the change in neutrophil populations, we examined the total numbers of neutrophils and their progenitors in the bone marrow (BM). Symptomatic infection in WT mice was accompanied by decreases in common myeloid progenitors CMPs as well as their derivative granulocyte-monocyte progenitors (GMPs) and megakaryocyte-erythrocyte progenitors (MEPs) (**Fig. 6A**). The size and ratio of these populations remained largely unaffected by infection in asymptomatic TCRα-deficient mice with only CMPs exhibiting a slight reduction (**Fig. S8A**). By contrast CMP numbers were reduced by 3-fold in infected WT mice compared to naïve WT controls. Across the two genotypes, the number of CMPs and MEPs was also reduced in infected WT mice in comparison to TCRα-deficient mice, whereas there was no difference between GMPs amongst the two groups. This was an important observation because there was a dramatic decrease in Ly6G^high^ neutrophils in the BM and the spleen of infected WT mice (**Fig. 6B**). Ly6G^low^ neutrophils were only reduced by approximately 50% which resulted in these cells becoming the predominant neutrophil population in the BM and in the periphery. By contrast, TCRα-deficient mice maintained a substantial Ly6G^high^ population and their Ly6G^low^ neutrophil numbers remained unchanged in the BM.

**Figure 6.**
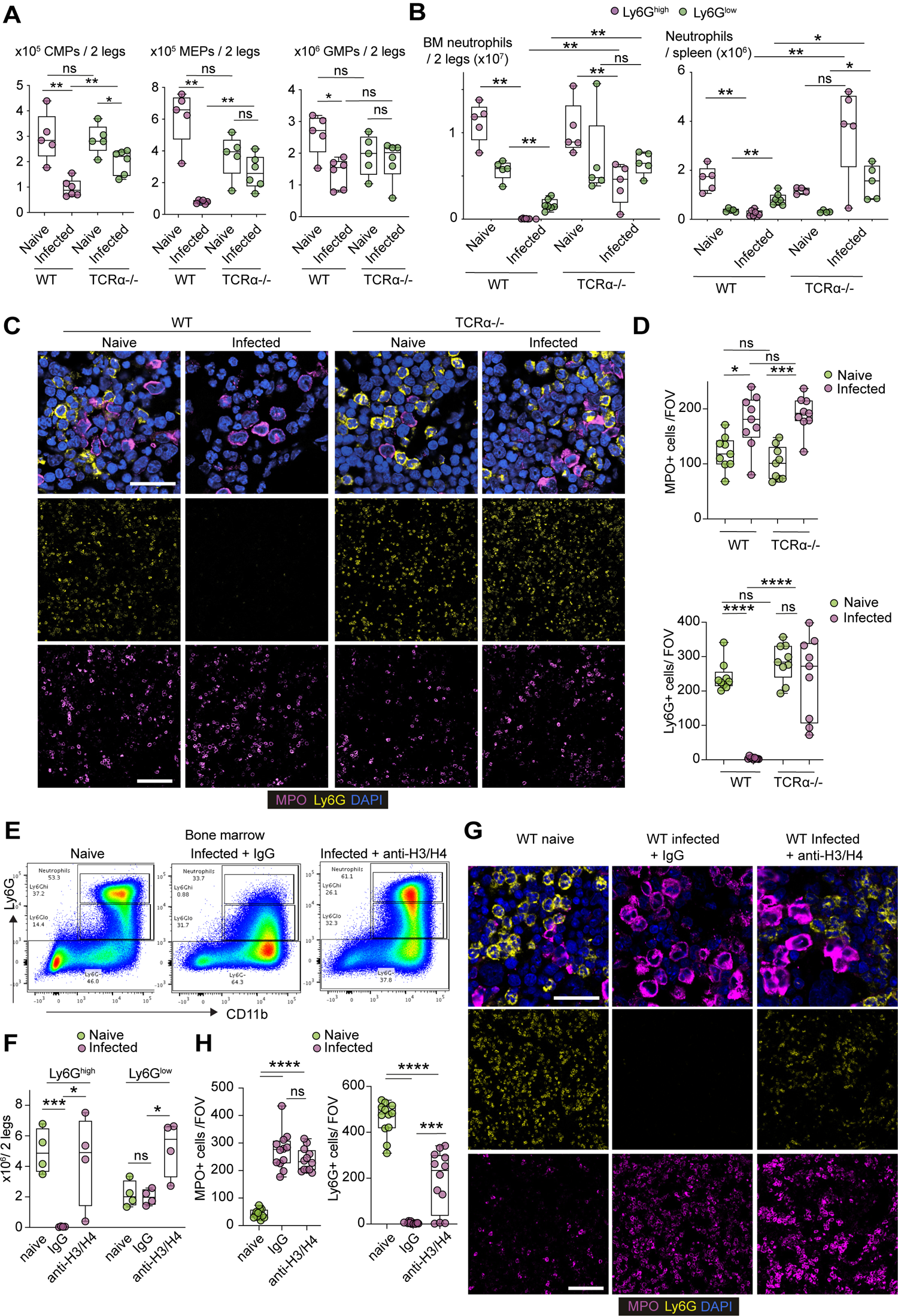
Histones deplete mature Ly6G^high^ neutrophils in the bone marrow. A. Flow cytometric analysis of myelopoietic progenitors and mature Ly6G^high^ neutrophils in the bone marrows of naive and infected WT and TCRα-deficient animals. Total numbers of common myeloid progenitors (CMP), megakaryocyte-erythrocyte progenitors (MEP) and granulocyte-monocyte progenitors (GMP) are shown. HSCs: Lineage (CD3, CD4, CD8, GR1, B220 and TER119)-c-KIT+ Sca-1-. MEPs: CD16/32 ^low/-^ CMPs: CD16/32^intermediate^, CD34+GMPs: CD16/32^high^. B. Total Ly6G^low^ and Ly6G^high^ neutrophils in the bone marrow (BM) and the spleens of naïve and infected WT and TCRα-deficient mice. 5 mice per condition were analysed. C. Immunofluorescence micrographs from the BM of naïve and infected WT and TCRα-deficient mice 3 days post infection stained for MPO, Ly6G and DAPI. D. Number of MPO+ and Ly6G+ neutrophils per field of view (FOV). Three FOV in 3 mice per condition were analysed. E. Representative flow cytometry plots of neutrophils in the bone marrow of naïve and infected WT mice pre-treated with a control IgG antibody (rabbit IgG) or antibodies against histones H3 and H4, 3 days post infection. F. Total Ly6G^low^ and Ly6G^high^ neutrophils in measured in (E). G. Immunofluorescence micrographs from the BM of naïve and infected WT mice pre-treated with an isotype IgG or antibodies against histones H3 and H4, 3 days post infection stained for MPO, Ly6G and DAPI. H. Quantification of the number of mature (Ly6G^+^) and immature (MPO^+^) cells per field of view (FOV) in (G) Three FOVs per mouse and four separate mice were analysed. Statistical analysis by unpaired Mann-Whitney t-test (* p<0.05, ** p<0.01, *** p<0.001, **** p<0.0001).

We took advantage in the relative difference between MPO and Ly6G expression levels in Ly6G^low^ and Ly6G^high^ neutrophils as determined by RNAseq (**Fig. 5C**), in order to distinguish these cells by immunofluorescence microscopy. Consistently, the MPO signal in immature Ly6G^low^ cells in the BM was high compared to mature Ly6G^high^ neutrophils which enabled the specific detection of immature neutrophils by immunofluorescence microscopy. Similarly, Ly6G staining detected specifically mature Ly6G^high^ neutrophils. As in the flow cytometry analysis, Ly6G^high^ neutrophils were completely absent from the BM of infected WT mice but were still present in infected TCRα-deficient BM (**Fig. 6C and 6D**). These findings indicated that neutrophil population changes occurred predominately at the late maturation stage, rather than during early differentiation, with profound loss of mature Ly6G^high^ neutrophils. Importantly, the infection impacted predominately mature neutrophil populations and to a much lesser degree immature granulocytes and their progenitors, since the BM of symptomatic WT and asymptomatic TCRα-deficient contained relatively equal numbers of GMPs and immature neutrophils.

To assess the role of histones in this process, we also examined the impact of histone-blocking antibodies on BM neutrophil populations. A cocktail of anti-H3 and anti-H4 antibodies maintained a significant presence of mature Ly6G^high^ neutrophils in the BM detected by flow cytometry (**Fig. 6E and 6F**) and by microscopy (**Fig. 6G and 6H**), indicating that histones played an important role in eliminating mature neutrophils from the BM.

### Histones and G-CSF disbalance neutrophil populations by modulating neutrophil lifespan

To investigate how these signals could impact the abundance of distinct neutrophil populations, we sorted Ly6G^low^ and Ly6G^high^ neutrophils from the BM of naïve WT mice as well as Ly6G^low^ from infected symptomatic WT mice and incubated them with varying concentrations of recombinant G-CSF and histone H3 alone or in combination. We monitored cell death by time-lapse immunofluorescence microscopy for 1300 min (**Fig. 7A**). We then fitted the cell death curves over time and obtained the half-life of these populations. Untreated mature Ly6G^high^ neutrophil populations from naïve mice had a slightly lower lifespan than Ly6G^low^ cells from naïve mice, whereas Ly6G^low^ cells from infected mice exhibited the longest lifespan (**Fig. 7B, S8B and S8C**). G-CSF treatment potently reduced the lifespan of Ly6G^high^ neutrophils but minimally affected naive Ly6G^low^ cells and extended the lifespan of Ly6G^low^ cells from infected mice, resulting in a large difference between the lifespans of Ly6G^high^ and Ly6G^low^ cells, particularly the ones isolated from infected symptomatic mice. Recombinant histone H3 treatment alone also decreased the lifespan of all three populations equally. A combination of modest concentrations of histone H3 (500nM) and G-CSF (1ng/mL) led to the largest decreased in mature neutrophil lifespan and the largest difference compared to the lifespan of Ly6G^low^ cells from infected mice. These data suggested that these factors shifted neutrophil populations disproportionately in favour of immature neutrophils by selectively shortening the lifespan of mature cells.

**Figure 7.**
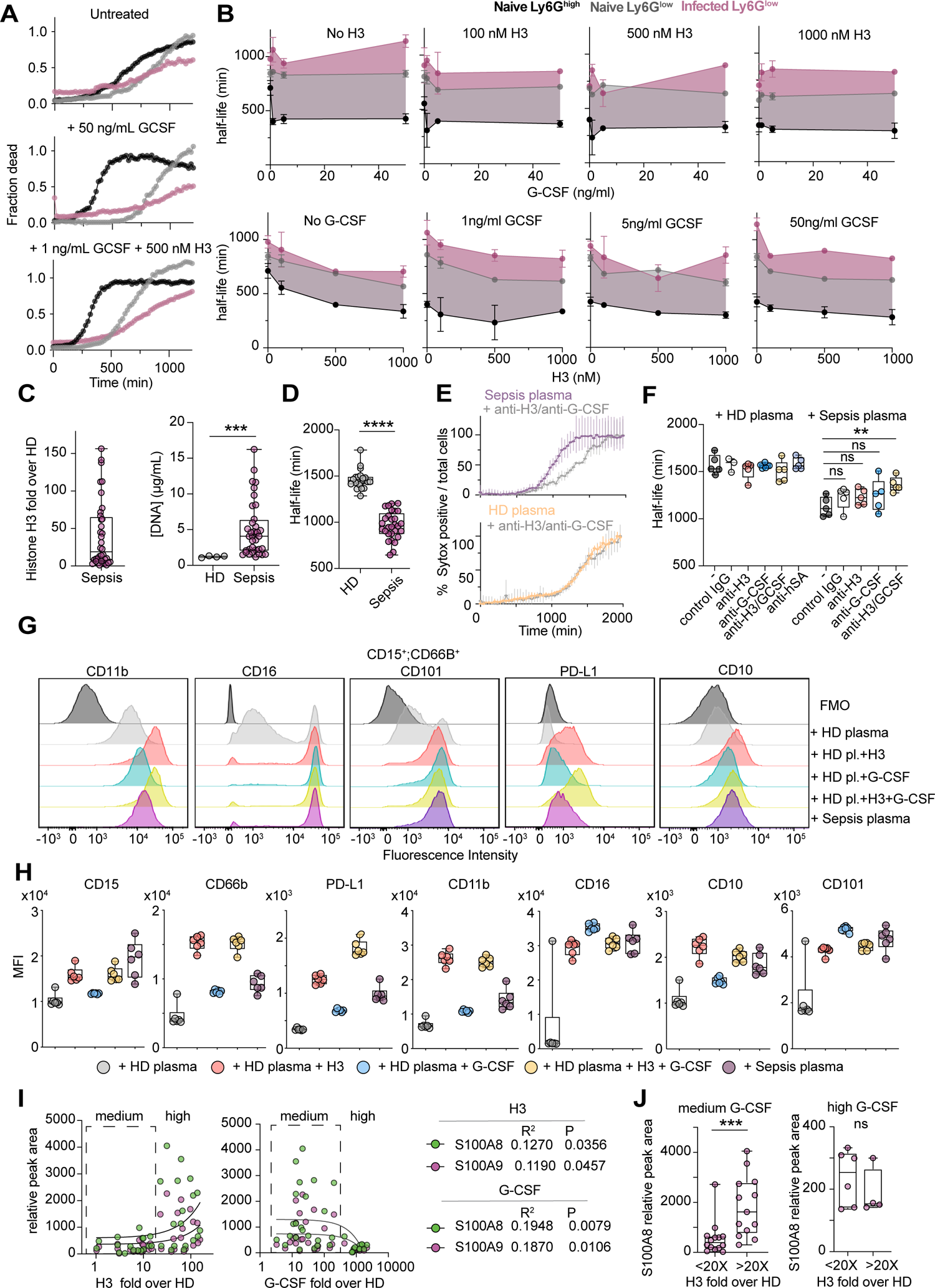
Histones and G-CSF regulate neutrophil lifespan A. Representative individual cell death curves over time of Ly6G^high^ (black) and Ly6G^low^ (grey) BM neutrophils sorted from naïve WT mice and Ly6G^low^ cells (purple) sorted from the BM of symptomatic infected WT mice populations in the absence of any stimuli (top) in the presence of 50 ng/mL G-CSF (middle), or in the presence of 1 ng/mL G-CSF and 500nM recombinant histone H3 (bottom). Data are representative of Representative of two independent experiments. B. Neutrophil half-lives measured from 1300 min-long cell death timecourse experiments in the presence of different concentrations of recombinant histone H3 and G-CSF alone or in combination. The difference in half-life between of Ly6G^low^ and Ly6G^high^ neutrophils is depicted by a coloured window mapped by the half-life curves using the corresponding color (grey for naive Ly6G^low^ and purple for infected Ly6G^low^). Each point is the average half-life from three separate field measurements. C. Fold increase in the levels of histone H3 in the plasma from human patients with bacterial sepsis over healthy donors (HD), quantified from Western immunoblots and expressed as a fraction over the average histone H3 signal from 4 HDs (left). DNA concentrations in the plasma of HDs and sepsis patients (right). D. Half-life of primary blood neutrophils from healthy donors incubated with either plasmas from different healthy donors (grey) or from sepsis patients (purple). Representative of two independent experiments. E. Representative cell death curves over 2000 min of healthy donor neutrophils incubated with HD (lower) or sepsis patient plasma (upper) in the presence of a control IgG antibody or antibodies against histone H3 and G-CSF. F. Quantification of the half-life of HD neutrophil populations incubated with either HD plasma or sepsis plasma alone or in the presence of either histone H3 blocking or G-CSF blocking antibodies separately or together. G. Representative histograms of flow cytometry analysis CD11b, CD16, CD101, PD-L1, and CD10 surface markers on healthy donor CD15+CD66b+ neutrophils incubated with HD or sepsis patient plasma alone or HD plasma supplemented with histone H3 or G-CSF or in combination for 1000 min. H. Quantification of the mean fluorescence intensity (MFI) of CD15, CD66b, PD-L1, CD11b, CD16, CD10, CD101 surface markers in healthy donor neutrophils incubated with HD or sepsis patient plasma alone or HD plasma supplemented with histone H3 or G-CSF or in combination and analysed via flow cytometry. Each plot shows includes 6 technical replicates, corresponding to different plasma donors. Data are representative of two independent experiments. I. Correlation between the fold-increase in plasma histone H3 (left plot) or G-CSF (right plot) and the relative abundance of S100A8 (green) and S100A9 (purple) detected by mass spectrometry in the plasma of sepsis patients. Boxes depict medium and high histone H3 containing plasmas. Boxes depict medium histone H3 or G-CSF-containing plasmas against the high content plasmas. J. Concentration of S100A8 in medium (left panel) and high (right panel) G-CSF-containing plasmas grouped based on medium (<20X fold over HD) or high (>20X fold over HD) histone H3 plasma concentrations. Statistical analysis by unpaired Mann-Whitney t-test (* p<0.05, ** p<0.01, *** p<0.001, **** p<0.0001).

To investigate the clinical relevance of this mechanism in human sepsis we measured the levels of G-CSF and histones in the plasma of sepsis patients with early (<24 hours) and severe (norepinephrine requirement > 0.4μg/kg/min) bacterial septic shock (*37*). We examined a group of patients with bacterial infections because bacterial sepsis cases are more prominent than fungal-associated sepsis, and although the upstream mechanisms regulating the induction of these cues may vary, we sought to understand whether the pathogenic signals played a relevant role in bacterial sepsis. Most patients exhibited elevated histone H3 and DNA concentrations in their blood when compared to healthy donors (**Fig. 7C**). Therefore, we incubated primary human neutrophils from the blood of healthy volunteers with 3% sepsis patient plasma and measured their lifespan over a 2000 min time-course. Sepsis plasma reduced mature human neutrophil lifespan by approximately 30% (**Fig. 7D**). This decrease depended on G-CSF and histones since neutralising antibodies against histone H3 and G-CSF significantly blocked lifespan reduction (**Fig. 7E and 7F**). The long duration of these lifespans was inconsistent with the immediate activation of cell death as cells persisted for 9 hrs before gradually starting to die. By contrast apoptosis is a rapid process (*38*). We also observed changes in neutrophil surface markers analysed by flow cytometry after 1000 min of incubation with sepsis plasma, compared to neutrophils incubated with healthy plasma (**Fig. 7G**). Sepsis plasma, or HD plasma complemented with recombinant histone H3 and G-CSF, increased the surface expression of CD11b, CD15, CD16, CD66b, CD10, CD101 and PD-L1. These markers enable the identification of altered neutrophils in sepsis patients and may enable to monitor the degree of lifespan modulation (**Fig. 7H**).

To assess the impact of this mechanism on sepsis severity, we generated a proteomic dataset for these patient plasmas and performed and unbiased search for significant correlations of histones and G-CSF with changes in plasma protein abundance. Interestingly, we found that histone H3 correlated with increased concentrations of cell-free plasma S100A8/9, which are proteins expressed in the cytosol of neutrophils and are released upon neutrophil death. Moreover, elevated plasma S100A8/9 in sepsis patients is associated with increased risk for mortality (*38*). Our analysis indicated that S100A8/9 levels were elevated in patients with intermediated G-CSF plasma concentrations but were suppressed by very high G-CSG concentrations (**Fig. 7I**), likely to be due to improved emergency granulopoiesis which may counter the detrimental effects of G-CSF on neutrophil lifespan (*7, 39, 40*). To better understand the impact of both plasma histones and G-CSF, we separated patients into medium and high G-CSF and histone groups and compared their plasma S100A8/9 content. Consistently, there was a significant increase in S100A8/9 in patients with medium G-CSF and high histone H3 plasma concentrations (**Fig. 7J**). Therefore, cell-free histones and G-CSF alter neutrophil lifespan during sepsis in a manner that is consistent with a gradual reduction in mature neutrophil populations over the course of infection.

## Discussion

Our findings uncover a mechanism linking SIGNR1^+^ macrophages to T cell death and neutrophil dysfunction. Mature neutrophils control fungi captured by SIGNR1^+^ macrophages via MPO-derived ROS. Progressive fungal colonization of the MZ promotes T cell death and the release of extracellular chromatin that synergizes with hyphae to induce G-CSF in CD169^+^ MZ macrophages. Sustained G-CSF production and T cell-derived chromatin compromised antimicrobial function by eliminating competent neutrophils and leaving behind immature cells with a defective ROS burst. These findings suggest that the spleen MZ forms an internal anti-microbial barrier, and its disruption accelerates the onset of sepsis pathology. Hence, immune dysfunction in neutrophils and T cells is linked, with neutrophils attenuating T cell death via microbial control and T cell death promoting the release of pathogenic factors that deplete mature immunocompetent neutrophils.

The importance of the spleen in antifungal defence was recently highlighted by the increased susceptibility of splenectomised mice challenged systemically with *C. albicans* (*41*). Similarly, asplenic patients are at a high risk of bacterial and fungal infection (*42*). Our study is consistent with the notion that while a healthy spleen is essential for the effective clearance of circulating pathogens, overwhelming this line of defence is detrimental. Similarly, while SIGNR1^+^ and CD169^+^ macrophages play beneficial roles in clearing blood-borne bacterial and viral infections (*43–45*), retroviruses can hijack this process to infect T cells (*46*). Similarly, our results indicate that *C. albicans* can take advantage of SIGNR1-mediated capture to initiate a programme that disrupts neutrophil antimicrobial function. Whether this process applies to bacterial infections is unclear, particularly since MPO deficiency has a minor effect on bacterial sepsis caused by cecal ligation and puncture (*47*). However, our experiments with bacterial sepsis patient samples indicate that the downstream signals that influence neutrophil lifespan are relevant in systemic bacterial infections.

Our study sheds light on mechanisms regulating the release of circulating histones in sepsis. Neutrophils are not a major source of histones as shown previously by neutrophil depletion experiments (*48*) and confirmed by the upregulation of extracellular chromatin in infected MPO-deficient mice. Here, we report for the first time, a cell population whose depletion abrogates plasma chromatin release suggesting that dying T cells are the key regulators and most likely source of cell-free chromatin in the circulation. While the mechanisms that drive T cell death in the spleen remain undefined, this process may depend on signals from CD169^+^ macrophages as demonstrated in other models via the action of monocyte-derived Fas ligand (*17*). Consistently, the activation of caspase-8 and caspase-3, TUNEL staining, and chromatin fragmentation are all indicative of T cell apoptosis. However, death of other cell types may also promote histone release into the circulation as demonstrated by our macrophage depletion experiments with Clo-L in uninfected mice. Furthermore, our study indicates that endogenous circulating free chromatin is not sufficient to cause acute pathology in the absence of infection. The amount of histones released by Clo-L treatment which did not cause pathology, was superior to the of histone release upon infection. Prior studies that demonstrated direct lethality in response to recombinant histone injections resulted in blood concentrations that were likely to be 100-fold higher than in naïve asymptomatic clodronate-treated animals (*5*). Moreover, the immunomodulatory properties of circulating chromatin suggest that caution should be exercised when Clo-L or any cell death-promoting agents are employed as their impact on cell death can alter innate immune production.

Our data indicate that MZ macrophages are important contributors of cytokines during systemic infection. The signals required to trigger various cytokines differ. Extracellular chromatin is sufficient to induce IL-6 and IL-1β (*9*). However, the induction of G-CSF requires the simultaneous presence of histones and fungal signals that signify an uncontrolled infection with a cytotoxic impact. During homeostasis, splenic macrophages regulate granulopoiesis by monitoring the presence of apoptotic circulating neutrophils (*49*). We now show that macrophages also link the detection of systemic microbes and DAMPs to the alteration of neutrophil populations.

The selective loss of mature neutrophils has also been demonstrated in murine bacterial sepsis models and linked to TREML4 (*21*). However, the cell death inducing signals and the basis of the selectivity for mature over immature cells remained unclear. Our study demonstrates that histones and G-CSF regulate this process selectively and may be relevant in other conditions, given that alterations in the age of neutrophils influences neutrophil function (*50–52*). The ability of G-CSF to shorten the lifespan of mature neutrophils explains how G-CSF can play both beneficial and detrimental roles during systemic infection. Furthermore, these signals induce a cell-surface signature that enables these cells and lifespan-reducing mechanisms to be detected. The different impact these signals exert on the lifespan of mature and immature neutrophils is a paradigm that explains the selective depletion of specific neutrophil populations.

Theoretically, the loss of mature neutrophils should expand the size of hematopoietic niches in the bone marrow (*49, 53*). However, all the progenitor numbers were reduced which may account for modest decreases in immature cells. The decrease in progenitors could be caused by the shift towards a ROS-deficient population given that ROS production is required for the expansion of progenitors during emergency granulopoiesis (*54*). Moreover, the elevated G-CSF levels may increase mature neutrophil egress in WT mice but can’t alone explain the loss of mature cells from the BM as the mature neutrophil numbers were low in the periphery. In addition, S100A8/9 proteins have recently been implicated in the induction of pathogenic neutrophil subsets in severe COVID-19 infection, suggesting by release of these alarmins, neutrophil lifespan-reduction could further impact neutrophil populations and disease severity (*23*).

The transcriptional profile of Ly6G^low^ neutrophils in symptomatic WT mice suggests that they are immature cells with additional infection-associated transcriptional changes, as they exhibit a distinct signature compared to Ly6G^low^ neutrophils in asymptomatic TCRα-deficient mice. Consistently, while the effect of sepsis on neutrophil can be nearly fully recapitulated by G-CSF and Clo-L administration, infection promotes additional changes in mature and immature neutrophils.

The basis for the different sensitivity of lifespan-shortening in mature and immature neutrophils to G-CSF and histones is unclear. Several transcriptional adaptations in the Ly6G^low^ neutrophil population of symptomatic mice may be important for survival. The downregulation of Csf3r transcript may result from a negative feedback in response to hyper-activation of G-CSFR. Ly6G^low^ cells maintained their surface protein expression of G-CSFR but they remain resistant to the lifespan reduction effects of its ligand. The downregulation of Clec2d may provide another mechanism that reduces Ly6G^low^ neutrophil exposure to histones given that this receptor mediates the uptake of extracellular chromatin (*55*). Moreover, the downregulation of GSDMD and GSDME transcripts and the upregulation of the anti-apoptotic factor Bcl2 may render Ly6G^low^ neutrophils more resistant to cell death.

In addition, several changes in gene expression may compromise the antimicrobial function of Ly6G^low^ neutrophils in symptomatic mice. A number of important genes for antimicrobial defence appear to be downregulated in these cells, such as Dectin-1 which is critical for fungal recognition and IL-1β which promotes neutrophil recruitment and swarming (*56, 57*). The predominant Ly6G^low^ neutrophil population exhibited lower ROS production capacity despite the expression of all major genes involved in NADPH oxidase function. Interestingly, Ly6G^low^ neutrophils exhibited a higher expression of MPO transcript and protein compared to Ly6G^high^ neutrophils allowing these cells to be distinguished from mature neutrophils in the bone marrow by microscopy. Lower ROS production also renders neutrophils more proinflammatory in fungal and bacterial infections which may contribute to sepsis pathology (*57*). However, the abundance of NETs in the kidneys of symptomatic mice, suggests that NETosis is not disrupted and may be driven either by the remaining minor mature neutrophil population or immature neutrophils with sufficient ROS burst. On the contrary, anti-SIGNR1 treatment reduced NETosis possibly by direct control of fungi via ROS, which may further reduction in pathology.

The pathogenic dysregulation of neutrophil populations by G-CSF and circulating histones may be relevant in other conditions such as cancer. Tumour-derived G-CSF is required but not sufficient to upregulate a minor population of pro-metastatic, tumour-induced low-density granulocytes (*36, 58, 59*). Tumour-induced DAMPs may play a role in driving pro-tumorigenic neutrophil populations and may provide further mechanistic understanding for the therapeutic potential of DNase treatment (*60–63*). Consistently, in our experiments, Ly6G^low^ neutrophils from infected symptomatic mice have increased arginase I expression which inhibits T cell proliferation and promotes tumour growth. Moreover, neutrophils from healthy human donors upregulated PD-L1 when incubated with sepsis plasma or G-CSF and histones, indicating that these conditions could enhance their pro-tumorigenic capacity. A sub-population of neutrophils with upregulated PD-L1 has also been described in COVID-19 patients and thus this mechanism may drive altered neutrophil populations in severe COVID-19 patients, where circulating chromatin and hyperinflammation have also been reported (*64, 65*).

Our study demonstrates that immune dysfunction in T cells and neutrophils is linked and can be targeted by interfering with microbial sequestration in sensitive areas and by eliminating circulating chromatin. The presence of G-CSF and circulating histones in many conditions suggests that they may play a generalized role as drivers of immune dysfunction that can be targeted therapeutically through histone-blocking strategies.

## Acknowledgements

We thank Andrew Aswani for sourcing sepsis patient samples, George Kassiotis, Brigitta Stockinger and Ilaria Malanchi for kindly providing reagents and mice and the Francis Crick Institute core facilities. This research was funded in whole, or in part, by the Wellcome Trust (FC0010129, FC001134). For the purpose of Open Access, the author has applied a CC BY public copyright licence to any Author Accepted Manuscript version arising from this submission. This work was supported by the Francis Crick Institute which receives its core funding from the UK Medical Research Council (FC0010129, FC001134), Cancer Research UK (FC0010129, FC001134) and the Wellcome Trust (FC0010129, FC001134). I.V.A was funded by an EMBO LTF (ALTF 113-2019).

## Author Contributions

M.I. and D.H. designed conducted and interpreted experiments, I.V.A performed and analysed neutrophil lifespan experiments, M.T. imaged and analysed lymph, thymus and BM samples, N.D.V. conducted macrophage activation experiments, T.D.T conducted hyphal growth experiments. Q.H. processed samples and stained tissues, S.B. and R.G. sequenced and analysed transcriptomic data, S.D., K.S., C.B. provided human sepsis plasma. S.V., V.D. and M.R. performed mass spectrometric analysis of patient samples. V.P. directed the study, analysed data and wrote the manuscript.

## Materials and Methods

### Bioinformatics primary data access

The high throughput RNA sequencing expression profile data can be accessed at NCBI Gene Expression Omnibus (GSE160301) and the token (ydyjocqyjpczjgt).

### Informed consent and ethics for human samples

For the in vitro neutrophil experiments, peripheral blood was isolated from consenting healthy adult volunteers, according to approved protocols of the ethics board of the Francis Crick Institute and the Human Tissue act. Sepsis patient samples were provided by the Hannover Medical School approved the ethics committee under the study protocol (No. 2786-2015). Written informed consent was obtained from participants or authorized representatives. The study was performed in accordance with the ethical standards laid down in the 1964 Declaration of Helsinki and its later amendments. No funding specific to this project was received.

### Animals

All mice were bred and maintained in a pathogen free, 12-hour light-dark cycle environment. All experiments were conducted with age-matched and cage-controlled, 8 to 12-week-old female WT C57BL/6J, MPO-/-(B6.129X1-*Mpo^tm1Lus^*^/J^), TCRα-/-(B6.129-Tcra*^tm1Phi^*), Rag2^-/-^ (B6.129S6-Rag2^tm1Fwa^ N12), G-CSF-/-(Csf3tm1Ard) and FVB/NJ mice, according to local guidelines and UK Home Office regulations under the Animals Scientific Procedures Act 1986 (ASPA).

### Neutrophil isolation from peripheral human blood

Peripheral venous blood was collected into EDTA tubes, layered on Histopaque 1119 (Sigma-Aldrich) and centrifuged for 20 min at 800x g. The plasma, PBMC and neutrophil layers were collected and neutrophils were washed in Hyclone Hank’s Balanced Salt Solution (HBSS) without calcium, magnesium or phenol red (GE Healthcare) supplemented with 10mM HEPES (Invitrogen) 0.1% plasma and further fractionated on a discontinuous Percoll (GE Healthcare) gradient consisting of layers with densities of 1105 g/ml (85%), 1100 g/ml (80%), 1093 g/ml (75%), 1087 g/ml (70%), and 1081 g/ml (65%) by centrifugation for 20 min at 800x g. Neutrophil enriched layers were collected and washed.

### In vitro stimulation of human neutrophils for FACs analysis

We isolated 2×10^5^ neutrophils human peripheral blood and seeded them in a U-bottom 96-well plate in HyClone HBSS +Ca, +Mg, - Phenol red (GE Healthcare) supplemented with 10mM HEPES (Invitrogen) and 3% plasma from healthy or septic donors. In vitro stimulation was performed with 500nM of human recombinant Histone 3 (Cayman chemical) and 5ng/ml recombinant human G-CSF (BioLegend). The cells were then incubated for 16 hours at 37°C and 5%CO2. Cells were then washed, stained and fixed for flow cytometry analysis according to the steps described below.

### Time-lapse imaging and half-life quantification

5×10^5^ neutrophils human peripheral blood were seeded in a black 96-well plate in HyClone HBSS +Ca, +Mg, - Phenol red (GE Healthcare) supplemented with 10mM HEPES (Invitrogen) and 3% plasma from healthy or septic donors. For mouse experiments, 5×10^5^ flow cytometry sorted bone marrow neutrophils were incubated with Roswell Park Memorial Institute 1640 medium (RPMI; Thermo Fisher Scientific) supplemented with 10mM HEPES (Invitrogen) and 3% foetal calf serum (FCS; Sigma). To distinguish between live and dead cells, we stained with 5uM Hoechst (membrane permeable; Thermo Scientific) and 5uM Sytox-green (membrane impermeable; Invitrogen). The cells were imaged on a long-term time-lapse wide-field Nikon system in a temperature (37°C) and CO2 (5%) regulation chamber. Four fields of view were acquired per well every 30 mins for 32 hours using a 40x objective.

Quantification of number of Sytox positive cells over total cells were performed using intensity-based thresholding masks in Fiji and particles between 20-1200 µm were quantified. % Sytox positive/total cells were fit with a sigmoidal function using Prism 9 to obtain the neutrophil half-lives. In vitro stimulation was performed with 100nM, 500nM and 1uM of human recombinant Histone 3 (Cayman chemical) and 1ng/ml, 5ng/ml or 50ng/ml recombinant human G-CSF (BioLegend) or mouse G-CSF (BioLegend). The blocking of H3 and G-CSF was performed by preincubating the different 3% plasma with 0.5 µg/ml of anti-hH3, 1 µg/ml and of anti-hG-CSF or 1.5 µg/ml of anti-IgG at 37°C for 30 mins before adding to the neutrophils.

### Sepsis in vivo model

Wild-type *Candida albicans* (*C. albicans*, clinical isolate SC5314) was cultured overnight shaking at 37oC and subcultered to an optical density (A_600_) of 0.4-0.8 for 4 hours in yeast extract peptone dextrose (YEPD; Sigma) medium. Subcultures were examined for lack of hyphae, washed and resuspended in sterile phosphate-buffered saline (PBS) immediately prior to infection. Mice were injected intravenously with either 1×10^3^, 1×10^4^, 1×10^5^ or 5×10^5^ *C. albicans* yeast particles per mouse. The weight and rectal temperature of the mice were recorded prior to infection and daily over the course of infection to track health status. A body temperature below 32°C, a weight loss superior to 80% of initial weight accompanied by slow movement and non-responsiveness were considered collectively as septic shock and the humane endpoint for the mice. The mice were culled via cervical dislocation or by lethal dose of pentobarbital (600mg/kg) with mepivacaine hydrochloride (20 mg/ml).

### Neutrophil and macrophage depletion in vivo

Neutrophil depletion was achieved with intraperitoneal injection of 150μg anti-Ly6G Ab (BioXCell) or IgG isotype control (BioXCell) at day - 1 and day 0 (day of infection). Macrophage depletion was performed with intravenous administration of 1mg Clodronate liposomes or 1mg PBS liposomes as control (Liposoma) at 1 day prior to infection.

### Treatment with blocking and neutralizing antibodies

SIGNR1 blocking was achieved with intraperitoneal injection of 100μg anti-SIGNR1 Ab (BioXCell) or hamster IgG isotype control (BioXCell) one day before infection followed by intraperitoneal injections (100μg/mouse) every other day. G-CSF neutralization was achieved with intraperitoneal injection of 100μg a-G-CSF (eBioscience) or IgG2a isotype control (eBioscience) started 1-2 days post-infection and performed every day until termination of the experiment. Histone neutralisation experiments were performed via intraperitoneal injection with dialysed and combined a-Histone 3 and a-Histone 4 antibodies (Merck Millipore) or control polyclonal rabbit IgG (BioXCell), starting on D-1 (200μg/mouse) and daily afterwards (200μg H3 and 100μg H4

### Treatment with recombinant proteins and enzymes

Recombinant mouse GCSF (BioLegend) was injected intraperitoneally (125μg/mouse) one day prior(D-1) or on the day of infection (D0) as indicated. For degradation of circulating DNA in vivo, mice were treated with deoxyribonuclease I (DNase I) from bovine pancreas (Sigma, 2000 units/mouse) or PBS vehicle control, intraperitoneally one day prior to infection and daily until completion of the experiment.

### Adoptive transfer experiments

T cells were isolated from naïve spleens with the EasySep T cell isolation kit (STEMCELL Technologies). For each mouse 4×10^6^ naïve T cells were injected intravenously via the tail, 2 days prior to infection.

### Induction of peripheral Ly6G^low^ neutrophils in naïve mice

Mice were injected intravenously with 1mg of Clo-L or PBS-L (Liposoma) and the subsequent day intraperitoneally with 2.5μg rG-CSF (BioLegend) or vehicle (PBS). Analysis of neutrophil populations was performed two days after rG-CSF injection, according to the methods described in flow cytometric analysis.

### *Candida albicans* culture for in vitro experiments

Wild-type *C. albicans* was cultured overnight shaking at 37°C in YEPD and then the following day sub-cultured to an optical density (A_600_) of 0.4-0.8 in YEPD (for yeast) or Roswell Park Memorial Institute 1640 medium (RPMI; Thermo Fisher Scientific) (for hyphae) for 4 hours. Cultures were checked for lack of hyphae (for yeast preparations) or for complete germination into hyphae and were then washed in PBS and counted. For hyphal growth assay live fungi were used and for in vitro stimulation of BMDMs, fungi were inactivated for 15min at 98⁰C.

### Hyphal growth assay

Bone marrow neutrophils were collected from the femur and tibia of naïve WT C57BL/6J and MPO-deficient mice using the EasySep mouse neutrophil isolation kit (STEMCELL Technologies). 2×10^5^ neutrophils were plated in Hank’s Balanced Salt Solution (HBSS) without calcium, magnesium or phenol red (Fisher Scientific) supplemented with 3% murine plasma and 10 mM HEPES (Sigma) and left to settle for 45 minutes in the incubator at 37°C and 5% CO2. Subsequently, 5×10^4^ hyphal units were added and the samples were imaged overnight with time lapse microscopy on a Leica DMIRB microscope (20x objective) fitted with an incubation chamber at 37°C and 5% CO_2_. Image analysis and quantification of hyphal length was performed with Fiji/ImageJ version 2.0.0 software.

### In vitro BMDM stimulation

Bone marrow was collected from femurs and tibias of wild type mice and cultured in Iscove’s Modified Dulbecco’s Medium (IMDM) + 30% L929 conditioned media + 10% FBS+ 1% Penincillin/Streptomycin (P/S). After 6 days of culture, bone marrow derived macrophages (BMDMs) were detached with PBS + 2mM EDTA, counted and 3×10^5^ cells were seeded per well in 24-well plates in IMDM + 10%FBS + 1%P/S. The next day, cells were washed to fresh IMDM + 10%FBS + 1%P/S media and stimulated overnight with increasing concentrations of human histone H3 (Cayman, cat. 10263) and/or heat inactivated *C. albicans* hyphae or yeast at multiplicity of infection (MOI) 3. Culture supernatants of stimulated BMDMs were collected and spun at 300*x*g for 10min at 4C. Supernatants were analysed for cytokines via methods described below.

### Tissue fungal burden

Organs were homogenized with a tissue homogenizer in PBS and plated in various dilutions on Sabouraud dextrose agar plates with 100μg/ml streptomycin to prevent growth of bacteria. The plates were incubated at 37°C for 12-18 hours and colony forming units (CFUs) were counted.

### Cytokines

Organs were homogenized with a tissue homogenizer in lysis buffer: PBS containing 1x proteinase inhibitors (Sigma Aldrich) and 0.1% Nonidet P-40 (NP40; Sigma). Cytokine and chemokine analysis from organ lysates and plasma were performed with the mouse 36-plex ProcartaPlex cytokine/chemokine array kit (ThermoFisher Scientific), G-CSF mouse or human ProcartaPlex Simplex Kit (ThermoFisher Scientific), G-CSF ELISA kit (R&D Systems), IL-6 mouse ELISA kit (ThermoFisher Scientific) and IL-1β mouse ELISA kit (ThermoFisher Scientific), following manufacturer’s instructions. All ProcartaPlex samples were analysed using the Luminex Bio-Plex 200 system. ELISA plate results were measured with light absorbance at 450 nm using a plate reader (Fluostar Omega, BMG labtech).

### Histology and Immunofluorescence imaging

Freshly extracted organs were embedded in optimal cutting temperature (OCT) compound cryo-embedding media (VWR Chemicals BDH) and flash-frozen in a dry ice/100%ethanol slurry. Frozen sections (8μm thickness) were dried, fixed in 4% paraformaldehyde (PFA; Sigma) and permeabilized with 0.5% Triton X-100 in PBS. Nonspecific binding was blocked with 2% BSA (Sigma) and 2% donkey serum (Sigma) in PBS. Samples were then stained with 4′,6-diamidino-2-phenylindole dihydrochloride (DAPI; Life Technologies) and the following primary antibodies: anti-B220 (BioLegend), anti-Candida (Acris), anti-CD3 (BioLegend), anti-CD4 (BioLegend), anti-CD169 (BioLegend), anti-F4/80 (BioLegend), anti-Ly6G (BioLegend), anti-MARCO (BMA Biomedicals), anti-MPO (R&D Systems), anti-SIGNR1 (eBioscience), anti-TCR-β (eBioscience), anti-cleaved-caspase 3 (Cell Signaling) and anti-cleaved-caspase 8 (Cell Signaling). When required, fluorescently labelled secondary antibodies were used: donkey anti-rabbit IgG (Invitrogen) and goat anti-rat IgG (Invitrogen).

For terminal deoxynucleotidyl transferase dUTP nick end labeling (TUNEL), the Click-iT TUNEL Alexa Fluor 594 Imaging Assay kit (Invitrogen) was used. Cryosections(8μm) were dried, fixed in PFA and permeabilized with 0.5% Triton X-100 in PBS. The continuing steps were performed as stated by the manufacturer and afterwards the slides were stained with primary and secondary antibodies as indicated previously.

RNAscope in situ hybridization assay was performed on frozen sections (8μm) using RNAscope 2.5 LS probes for G-CSF and IL-6 (Mm-CSF3-C2, cat. 400918, Mm-IL6 cat. 315898-C1, ACD/Bio-Techne) using the automated Assay for Leica Systems and according to the manufacturer’s instructions. Subsequently, the slides were stained with primary and secondary antibodies following the methods described above.

Pan-caspase labelling was performed using the Poly Caspase Assay Kit (MyBioSource) according to the instructions provided by the manufacturer.

All stained tissue sections were mounted in ProLong Gold (Molecular Probes). Images were taken using the Leica TCS SP5 inverted confocal microscope (20x, 40x, 63x original magnification) and analysis was performed using Fiji/ImageJ version 2.0.0 software.

### Tissue dissociation and preparation of single cell suspension

Kidneys were chopped up and incubated in a digestion medium containing 0.2mg/ml Liberase TL (Roche) and 0.1mg/ml DNase I (Roche) for 20 minutes while shaking at 37°C. The digested kidney tissue was filtered and centrifuged in a Percoll (GE Healthcare) density gradient (40%/70%). Spleens were gently meshed in FACs buffer (PBS containing 3% FCS from Sigma) using a 40μM cell strainer to prepare single cell suspensions. Whole blood was centrifuged in Histopaque-1119 (Sigma) in order to isolate peripheral blood leukocytes. ACK (ammonium-chloride-potassium, GIBCO) lysing buffer was used in all tissue samples to eliminate remaining erythrocytes.

### Flow cytometric analysis and sorting

Single cell suspensions were stained for 25 mins at 4°C with LIVE/DEAD^TM^ fixable dead cell stains (Thermofisher Scientific) or DAPI (Invitrogen) and the following fluorescent antibodies: anti-CD3, anti-CD19, anti-CD11b, anti-Ly6G, anti-Ly6C, anti-C5aR1, anti-CD66b, anti-CD101, anti-PD-L1 (all from BioLegend), anti-G-CSFR primary (R&D), anti-rat secondary (Invitrogen), anti-CD15, anti-CD16, anti-CD10 (all from BD Biosciences) and MPO (Hycult Biotech). For intracellular stainings, FoxP3 transcription factor staining buffer kit (eBiosciences) was used following manufacturer’s instructions. The BD LSR Fortessa was used for acquisition of all samples and FlowJo software v10 for subsequent analysis.

For the Vβ T cell analysis, single cell suspensions were obtained as indicated above and the mouse Vβ TCR Screening Panel (BD Pharmingen) was used for staining, according to the instructions provided by the manufacturer.

For sorting single cell suspensions were obtained as described prior and enriched with the EasySep mouse neutrophil isolation kit (STEMCELL Technologies). This semi-pure cell suspension was stained with fluorescently labelled antibodies via the methods described above and sorted with a 70uM nozzle (RNAseq and Giemsa stainings) or a 100uM nozzle (in vitro culture experiments) on the BD FACSAria cell sorter. Cells were sorted in pure FCS.

### Cytospin and Giemsa staining

For neutrophil characterization by nuclear morphology, 10^5^-10^6^ sorted neutrophils were cytospun on slides, fixed in methanol and stained with freshly prepared and filtered Giemsa staining buffer, containing Giemsa R (Ral Diagnostics). All slides were mounted in dibutylphthalate polystyrene xylene (DPX) and imaged using the ZEISS Axio Observer Z1.

### ROS assay

Mouse spleens were gently meshed using a 40μM cell strainer and erythrocytes were lysed with ACK lysing buffer (GIBCO). Neutrophils were isolated via negative selection with the EasySep mouse neutrophil isolation kit (STEMCELL Technologies). A total of 1×10^5^ neutrophils were then incubated with the chemiluminescent probe luminol (Sigma) and horseradish peroxidase (HRP; Sigma) and stimulated with 100nM PMA; (Sigma). Reactive oxygen species (ROS) production is measured by HRP mediated oxidation of luminol, producing a chemiluminescent signal that is detected with an UV filter on a spectrophotometric microplate reader (Fluostar Omega, BMG labtech).

### Western blot analysis

Samples (protein extracts or plasma) were boiled in a sodium dodecyl sulphate (SDS) buffer containing dithiothreitol (DTT) and resolved by polyacrylamide gel electrophoresis (SDS-PAGE) on a Criterion TGX precast gel (Any-KD; Bio-Rad Laboratories). Proteins were then transferred to a polyvinylidene difluoride (PVDF) membrane (Bio-Rad Laboratories) via semi-dry transfer. Nonspecific binding was blocked with 5% bovine serum albumin (BSA; Fisher Scientific) in tris-buffered saline with 0.1% Tween 20 (TBS-T). The membranes were blotted with anti-histone 3 (Milipore) and detected with HRP–conjugated goat anti-rabbit (Thermo Scientific). Finally, the membranes were incubated with enhanced chemiluminescent substrate (ECL; Thermo Fisher Scientific) and imaged with a chemiluminescence imaging systems (Bio-Rad) or developed after exposure onto an X-ray film (Kodak) and digitally scanned.

### DNA quantitation

DNA in plasma was quantified using the Quant-iTTM PicoGreen dsDNA assay kit (Thermo Fisher Scientific) following the instructions provided by the manufacturer. The fluorescent signal (excitation at 488nm) was measured using a spectrophotometric microplate reader (Fluostar Omega, BMG labtech).

### Plasma sample preparation for proteomic analysis

24 (heparin-treated) plasma samples of WT naive, WT infected, TCRα -/- naive, TCRα -/- infected mice in equal replicates of six were used for proteomics analysis. The samples were randomised and plated in a 96-well plate (Eppendorf). The protocol used for protein/peptide extraction and proteomics analysis has been described in detail in Messner et al., 2020 (*66*). Briefly, 5μL of plasma was denatured in 50μl 8M Urea (Honeywell Research Chemicals), 100mM ammonium bicarbonate (ABC, Honeywell Research Chemicals) and reduced with 5μL of 50mM dithiothreitol (DTT, Sigma Aldrich) at 30°C for 1 hour. Alkylation with 5μL of 100mM iodoacetamide (IAA, Sigma Aldrich) followed, at 23°C for 30 minutes in the dark. Then the samples were diluted with 340μL of 100mM ABC and 220μL were added to trypsin solution (Promega) for protein digestion at a trypsin/protein ratio of 1/40 and incubated at 37°C overnight (17h). Quenching of digestion was done by the addition of 25μL of 10% v/v formic acid (FA, Thermo Fisher Scientific). Rounds of solid phase extraction clean-up steps were performed with the use of C18 96-well plates (BioPureSPE Macro 96-Well, 100mg PROTO C18, The Nest Group) as described previously in Messner et al. 2020 (*66*). Methanol (Fisher Chemicals), 50% v/v acetonitrile (ACN, Fisher Chemicals) or 0.1% v/v FA was used at each centrifugation step as required. After final elution, the collected peptide material was dried by a vacuum concentrator (Eppendorf Concentrator Plus) and redissolved in 50μl 0.1% v/v FA, to be processed by liquid chromatography-mass spectrometry.

### Liquid chromatography-mass spectrometry

1μg of protein digest (peptides) was injected and analysed on a nanoAcquity Liquid Chromatograph (Waters) coupled to a TripleTOF 6600 Mass Spectrometer (Sciex) at a flow-rate of 5µl/min. Separation was achieved using a Waters HSS T3 column (150mm x 300µm, 1.8µm particles) in 20-minute non-linear gradients starting with 3% B up to 40% B (Buffer A: 0.1% v/v FA; Buffer B: ACN / 0.1% v/v FA). A data independent acquisition (DIA/SWATH) method was used, with MS1 scan from m/z 400 to m/z 1250 and 50ms accumulation time followed by 40 MS2 scans of 35ms accumulation time with variable precursor isolation width covering the mass range from m/z 400 to m/z 1250. Ion source gas 1 (nebulizer gas), ion source gas 2 (heater gas) and curtain gas were set to 30, 15 and 25 respectively. The source temperature was set to 450°C and the ion spray voltage to 5500V. Injections of samples took place in a random order.

The library was generated with “gas-phase fractionation” methodology out of pooled peptide digest of all mouse plasma samples with the use of a LC-MS/MS method as mentioned before. 3μg of protein digest was separated with a 60-minute linear gradient (3% B to 40% B). Injections were performed at the mass ranges of: 400 – 500 m/z, 495 – 600 m/z, 595 – 700 m/z, 695 – 800 m/z, 795 – 900 m/z, 895 – 1000 m/z, 995 – 1100 m/z, 1095 – 1250 m/z. The precursor selection windows were 2 m/z (1 m/z overlap). DIA-NN 1.7.10 proteomics analysis software (*67*) was used for the library preparation with Mus musculus (mouse) UniProt (UniProt Consortium, 2019) isoform sequence database (UP000000589) to annotate the library.

For protein quantification, raw data acquired were processed with DIA-NN 1.7.10 with the “robust LC (high precision)” mode with MS2, MS1 and scan window size set to 20ppm, 12ppm and 8 respectively.

### Preparation of samples for high throughput RNA sequencing

Spleen leukocytes were extracted via methods described in the flow cytometry section. Following homogenization, samples were enriched using the EasySep mouse neutrophil isolation kit (STEMCELL Technologies) and subsequently stained with antibodies and isolated via flow cytometric-sorting. RNA extraction was performed with the RNeasy mini kit (Qiagen) and the libraries were prepared with the Nugen cDNA synthesis kit.

### Bioinformatics

Sequencing was performed on an Illumina HiSeq 4000 machine. The ‘Trim Galore!’ utility version 0.4.2 was used to remove sequencing adaptors and to quality trim individual reads with the q-parameter set to 20 (1). Then sequencing reads were aligned to the mouse genome and transcriptome (Ensembl GRCm38 release-89) using RSEM version 1.3.0 in conjunction with the STAR aligner version 2.5.2(*68, 69*). Sequencing quality of individual samples was assessed using FASTQC version 0.11.5 and RNA-SeQC version 1.1.8(*70*). Differential gene expression was determined using the R-bioconductor package DESeq2 version 1.24.0(*71*). Gene set enrichment analysis (GSEA) was conducted as described in(*72*).

## Statistical analysis

Single comparison statistical significance was assessed by an unpaired, two-tailed Mann-Whitney t-test. Log-Rank Mantel-Cox test was applied for analysis in survival experiments. Statistical analysis was performed on GraphPad Prism software. P values of less than 0.05 were considered significant: n.s. p ≥ 0.05, * p ≤ 0.05, ** p ≤ 0.01, *** p ≤ 0.001.

---

**Figure S1.**
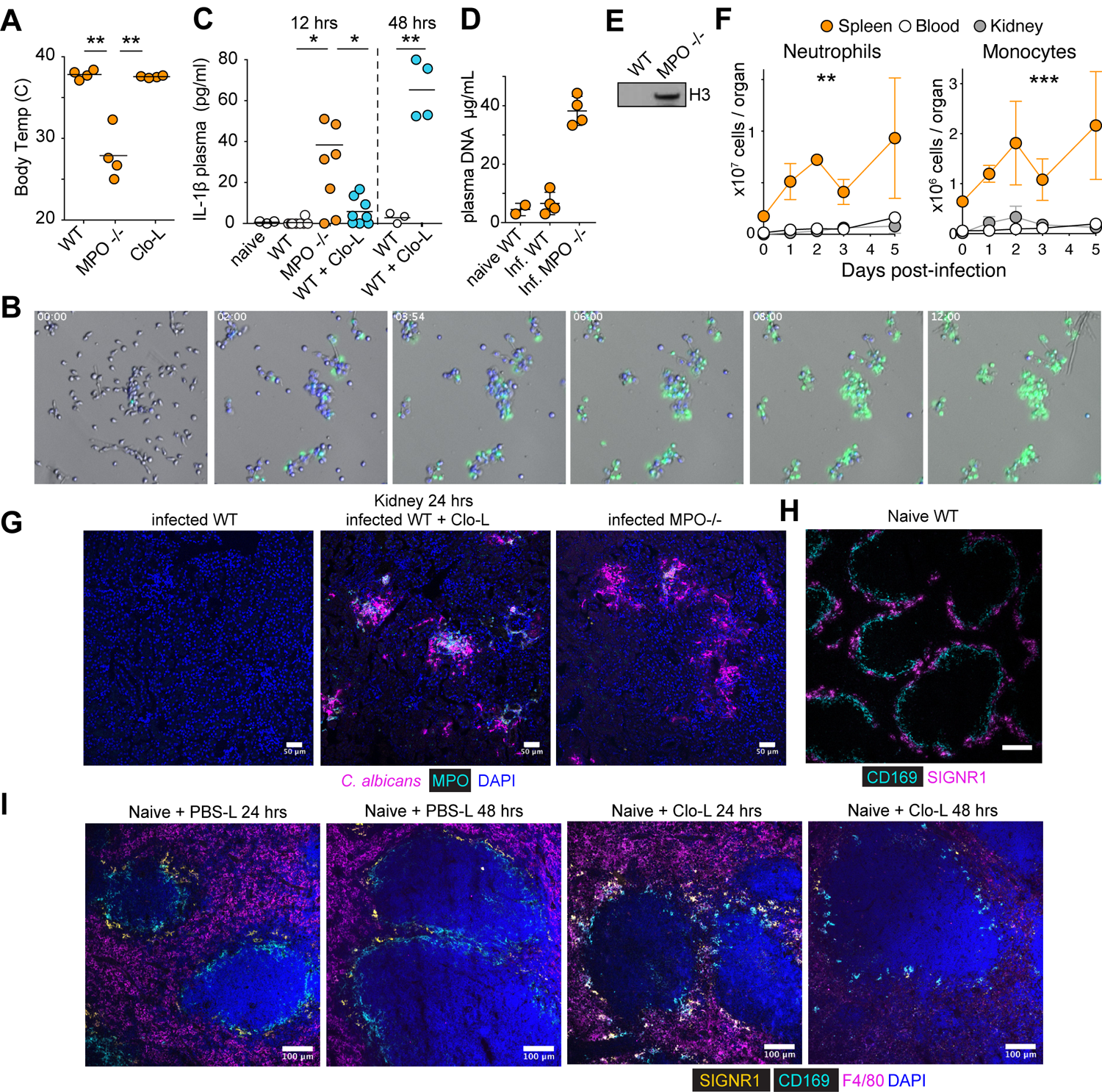
Effect of MPO deficiency on fungal colonization. A. Body temperature of WT mice or WT mice pre-treated with clodronate liposomes (Clo-L), or MPO deficient mice, infected intravenously with 1×10^5^ WT *C. albicans*, 12 hrs post-infection. B. Timelapse microscopy of *C. albicans* hyphae cultured with WT bone marrow derived murine neutrophils in the presence of Hoechst (blue) and Sytox Green (green) spanning time 0 to 12 hrs. Images were obtained every 2 min intervals. Representative images at T0, 2, 4, 6, 8, 12 hrs. C. IL-1β concentrations in the plasma of naïve WT mice, or WT, MPO-deficient mice or WT mice treated with Clo-L and infected intravenously with 1×10^5^ WT *C. albicans*, 12 hrs and 48 hrs post-infection. D. Plasma DNA concentrations from naïve WT and infected WT and MPO deficient animals 1 day post-infection. E. Representative Western immunoblot of histone H3 content in the WT and MPO-deficient mice infected with 1×10^5^ WT *C. albicans*, 12 hrs post-infection. F. Total number of neutorphils in the blood, spleen and kidneys of WT mice infected intravenously with 1×10^5^ WT *C. albicans* at the indicated day post-infection analyzed by flow cytometry. G-I Representative immunofluorescence confocal micrographs from: G. kidneys from WT, MPO-deficient mice or WT mice treated with Clo-L and infected intravenously with 1×10^5^ WT *C. albicans*. Stained for *C. albicans*, MPO and DAPI. Scale bars: 50 μm. H. spleen of a naïve WT mouse, stained for CD169 and SIGNR1. Scale bars: 100 μm. I. WT spleens 24 hrs and 48 hrs after administration of PBS liposomes (PBS-L) or clodronate liposomes (Clo-L) stained for CD169, SIGNR1, F4/80 and DAPI. Scale bars: 100 μm. Statistical analysis by unpaired Mann-Whitney t-test (* p<0.05, ** p<0.01, *** p<0.001, **** p<0.0001).

**Figure S2.**
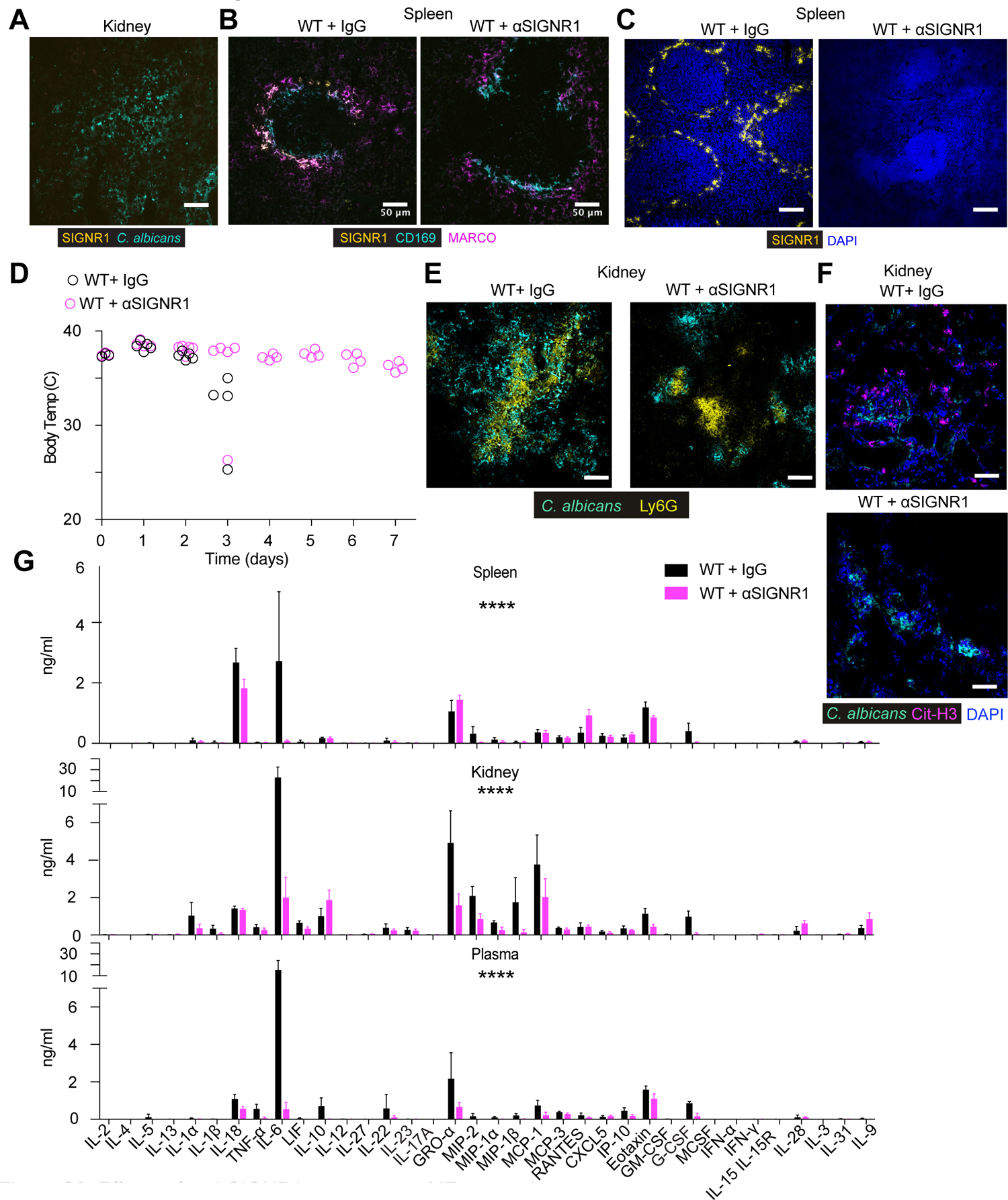
Effects of anti-SIGNR1 treatment on MZ macrophages. A. Representative immunofluorescence confocal micrograph from the kidney of symptomatic WT mice 3 days post-in-fection, stained for *C. albicans* and SIGNR1. Scale bars: 50 μm. Sections are representative of three mice analysed individually in at least 2 experimental replicates. B. Spleens of infected WT mice pre-treated with control antibody or an antibody against SIGNR1, 3 days post-infection, stained for SIGNR1, MARCO, CD169 and DAPI. Scale bars: 50 μm. C. spleen from a naïve WT mouse, treated with control IgG and anti-SIGNR1 antibody stained for SIGNR1 and DAPI. Scale bars: 50 μm. D. Body temperatures of WT mice pre-treated with either control antibody (black open circles) or an anti-SIGNR1 blocking antibody (magenta open circles) and infected intravenously with 5×10^5^ WT *C. albicans*. E. Micrographs of kidneys 3 days post-infection from WT mice pre-treated with either control antibody or an anti-SIGNR1 blocking antibody and infected intravenously with 5×10^5^ WT *C. albicans*. Stained for *C. albicans* and Ly6G. Scale bars: 50 μm F. Kidney samples in (E) stained for cit-H3, *C. albicans* and DAPI. G. Cytokines and chemokines from the blood, spleens and kidneys in WT mice pre-treated with either control antibody or an anti-SIGNR1 blocking antibody and infected intravenously with 5×10^5^ WT *C. albicans,* 3 days post infection. Statistical analysis by two-way Anova.

**Figure S3.**
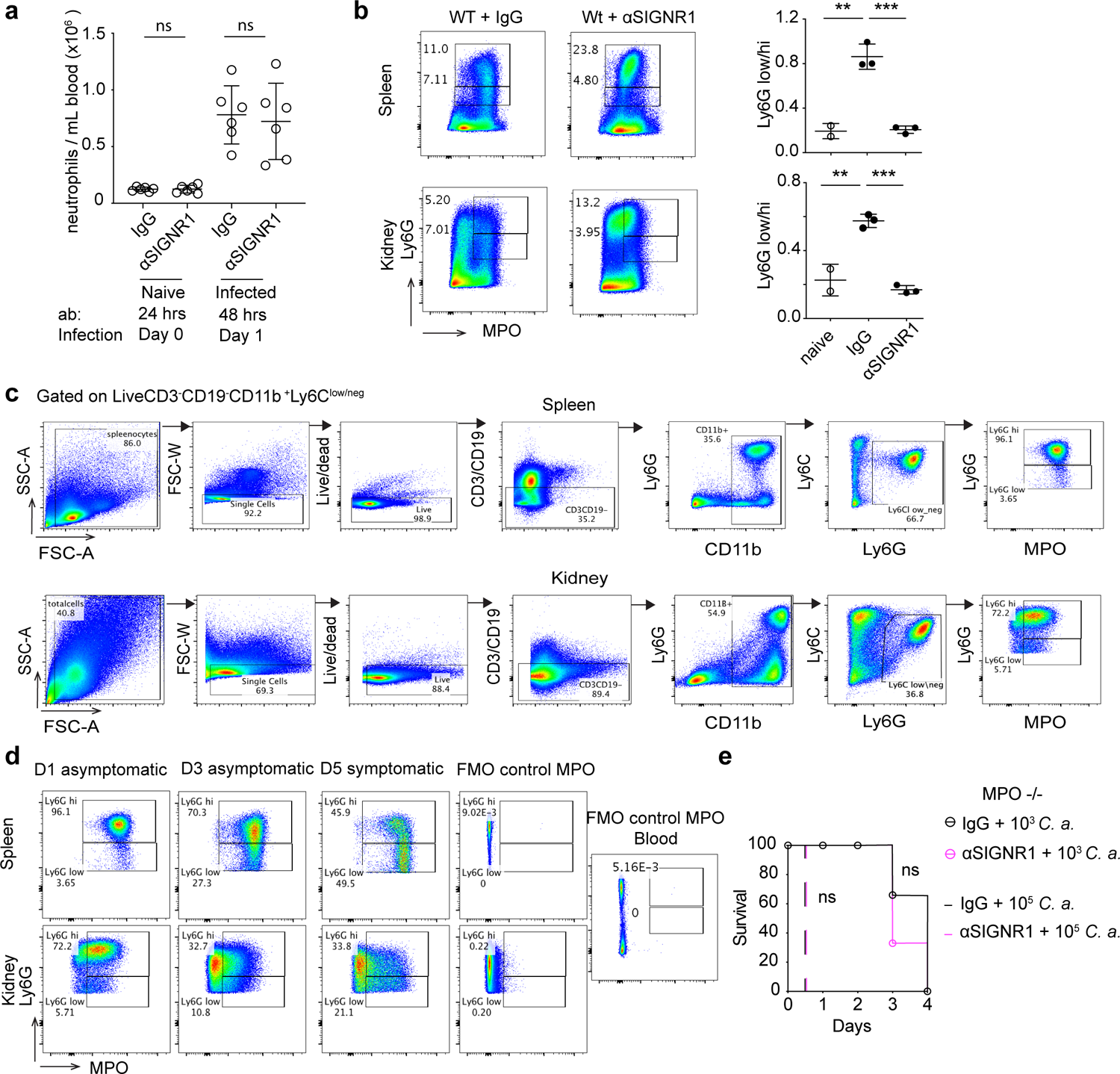
Effects of anti-SIGNR1 treatment on neutrophil populations. (A-B) Analysis of samples from WT mice pre-treated with either control antibody or an anti-SIGNR1 blocking antibody and infected intravenously with 5×10^5^ WT *C. albicans*. N=3-6 mice per group and at least 3 independent experiments. A. Total number of neutrophils per ml of blood in naïve mice on the day of infection (24 hrs after antibody administration) and infected mice 24 hrs post infection (48 hrs after antibody administration). B. Representative dot plots (left) and Ly6G^low^/Ly6G^high^ ratios (right) of flow cytometric analysed neutrophils in kidneys and spleens of infected mice pre-treated with IgG control or anti-SIGNR1 antibodies, 72 hrs post infection and stained for intracellular MPO and surface Ly6G expression. C. Gating strategy for flow cytometric analysis of neutrophils in the spleen and kindey. D. Flow cytometric analysis of neutrophils in the spleens and kidneys of WT infected mice collected at 24 hrs, 72 hrs and 5 days post-infection, in an experiment where WT mice started developing symptoms 5 days post-infection. E. Survival of MPO-deficient mice pre-treated with either control antibody or an anti-SIGNR1 blocking antibody and infected intravenously with 1×10^3^ or 1×10^5^ WT *C. albicans*. N=3-5 mice per group. Statistical analysis by unpaired Mann-Whitney t-test for single comparison and Log-rank (Mantel-Cox) test for survival analysis (ns >0.05, * p<0.05, ** p<0.01, *** p<0.001, **** p<0.0001).

**Figure S4.**
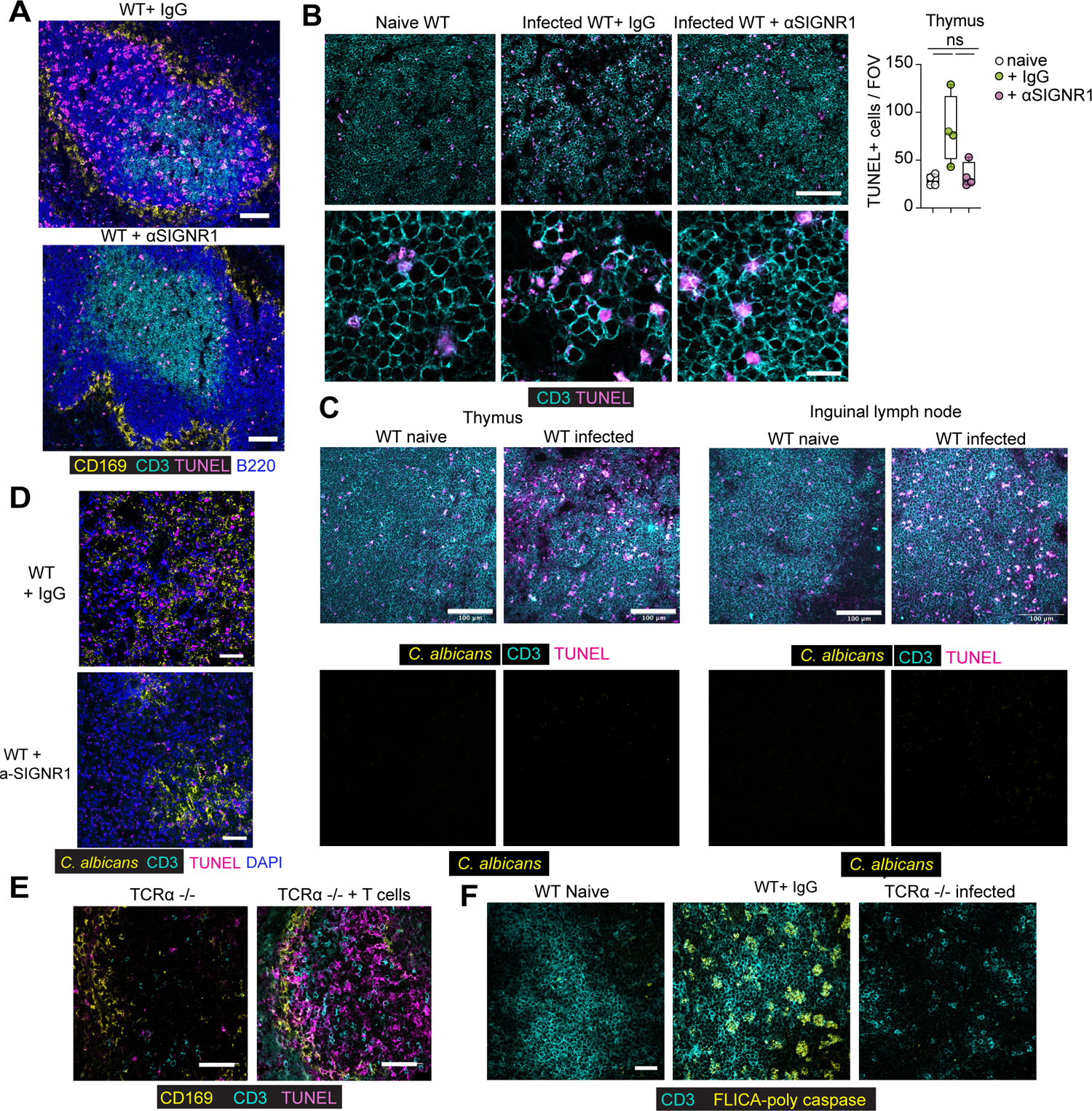
Effects of anti-SIGNR1 treatment on cell death in the kidney and lymph nodes. A. Immunofluorescence micrographs from spleen of WT mice pre-treated control antibody and infected with 5×10^5^ WT *C. albicans*, 3 days post-infection and stained for double stranded DNA breaks (TUNEL), CD3 CD169 and B220. Scale bars: 50 μm B. Immunofluorescence micrographs and quantification of thymus from WT mice, either naïve or infected with 5×10^5^ WT *C. albicans* and treated with control IgG or anti-SIGNR1 antibodies, 72 hrs post-infection and stained for TUNEL and CD3. Scale bars: 100 μm (upper row) and 25 μm (lower row). C. Immunofluorescence confocal micrographs of thymus and inguinal lymph nodes from WT mice, either naïve or infected with 5×10^5^ WT *C. albicans*, 72 hrs post-infection and stained for double stranded DNA breaks (TUNEL), CD3 and *C. albicans*. Scale bar: 100 μm. D. Immunofluorescence micrographs of kidneys from WT mice pre-treated with either control antibody or an anti-SIGNR1 blocking antibody and infected intravenously with 5×10^5^ WT *C. albicans*, at 72 hrs post-infection. Staining for *C. albicans*, CD3, TUNEL, DAPI. Scale bar: 50 μm. E. Immunofluorescence micrographs form the spleens of TCRα-deficient mice alone or after adoptive T cell transfer and infected with 5×10^5^ WT *C. albicans*, 72 hrs post-infection. F. Immunofluorescence micrographs from the spleens of naive or infected WT or TCRα-deficient mice infected with 5×10^5^ WT *C. albicans*, 3 days post-infection and stained for CD3 and poly-caspase enzyme activity performed with a FLICA poly-caspase activity assay. Scale bars: 50μm. Statistical analyses by unpaired Mann-Whitney t-test (* p<0.05, ** p<0.01, *** p<0.001, **** p<0.0001).

**Figure S5.**
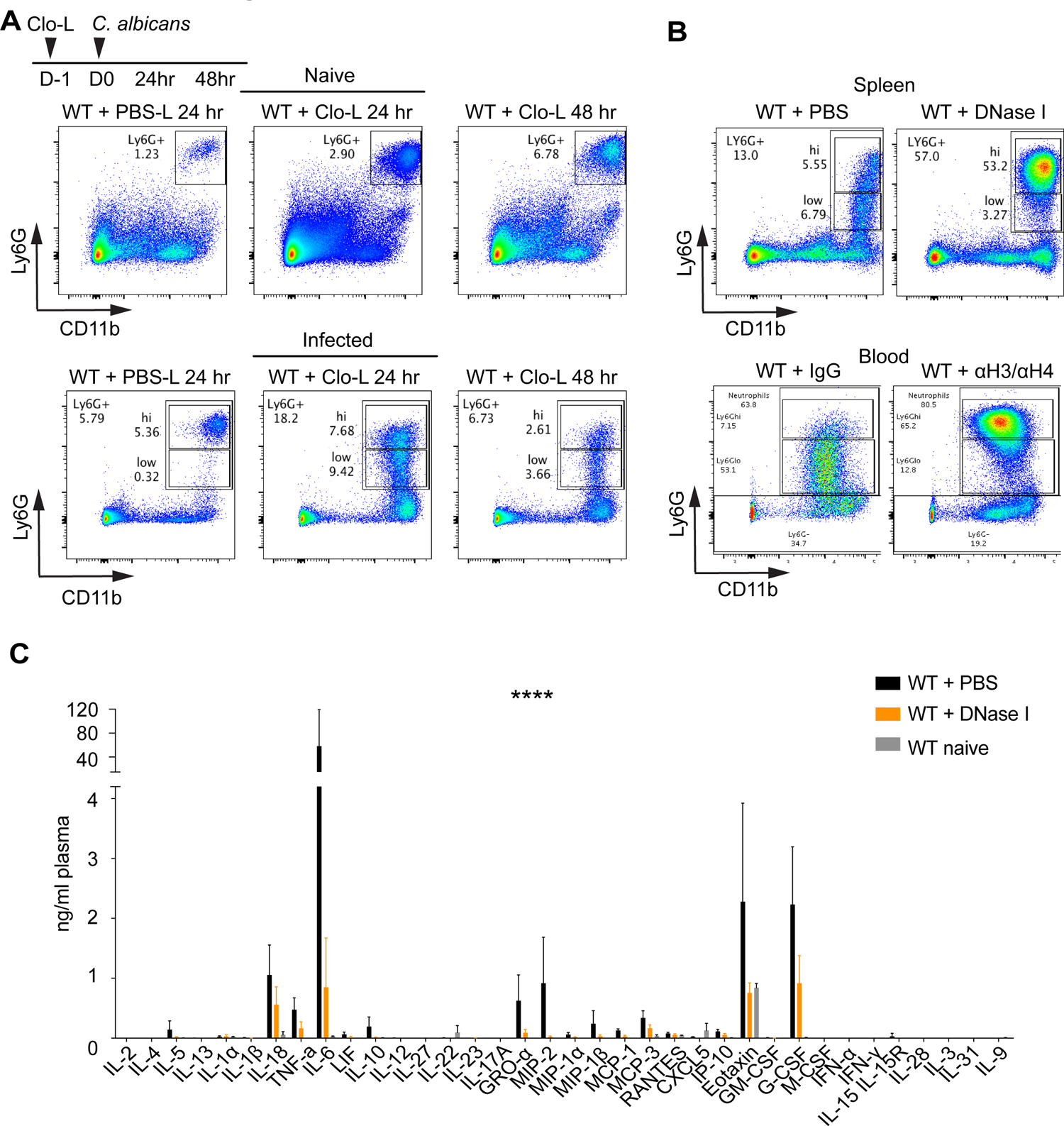
G-CSF and cell death-derived histones alter neutrophil populations. A. Representative flow cytometry plots depicting Ly6G^low^/Ly6G^high^ ratios in the spleens of WT mice pre-treated with PBS-L or Clo-L, either naïve or infected with 1×10^5^ WT *C. albicans*, 24 hrs and 48 hrs post-infection. B. Representative flow cytometry plots depicting Ly6G^low^/Ly6G^high^ ratios in the spleens and blood of WT mice pre-treated with either DNase I or PBS or control IgG or anti-H3/anti-H4 antibodies and infected with 5×10^5^ *C. albicans*, 3 days post-infection. C. Cytokines and chemokines measured by multiplex immunoassay in the plasma of WT mice pre-treated with DNase I or PBS and infected with 5×10^5^ *C. albicans*, 48 hrs post-infection. Statistical analysis by two-way Anova (* p<0.05, ** p<0.01, *** p<0.001, **** p<0.0001).

**Figure S6.**
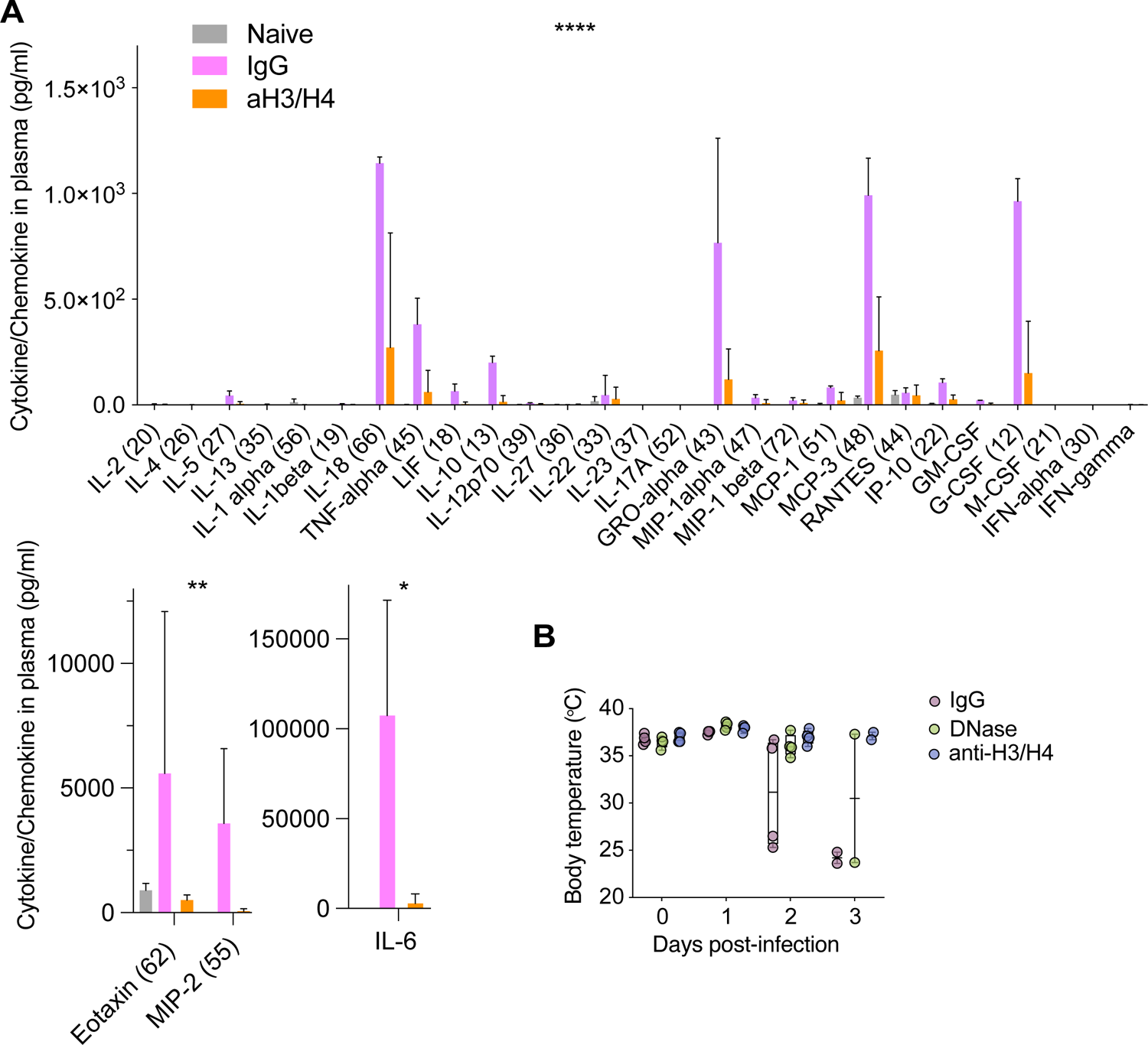
G-CSF and cell death-derived histones alter neutrophil populations. A. Cytokines and chemokines measured by multiplex immunoassay in the plasma of naive WT mice or pre-treated with IgG or anti-H3/anti-H4 antibodies and infected with 5×10^5^ *C. albicans*, 48 hrs post-infection. B. Body temperatures of infected WT mice treated with either an isotype control antibody (IgG), DNase I, or antibodies against histone H3 and H4 from experiment in Fig. 4J and Fig. 6E-H. Statistical analysis by two-way Avova and unpaired Mann-Whitney t-test for single comparison (IL-6) (* p<0.05, ** p<0.01, *** p<0.001, **** p<0.0001).

**Figure S7.**
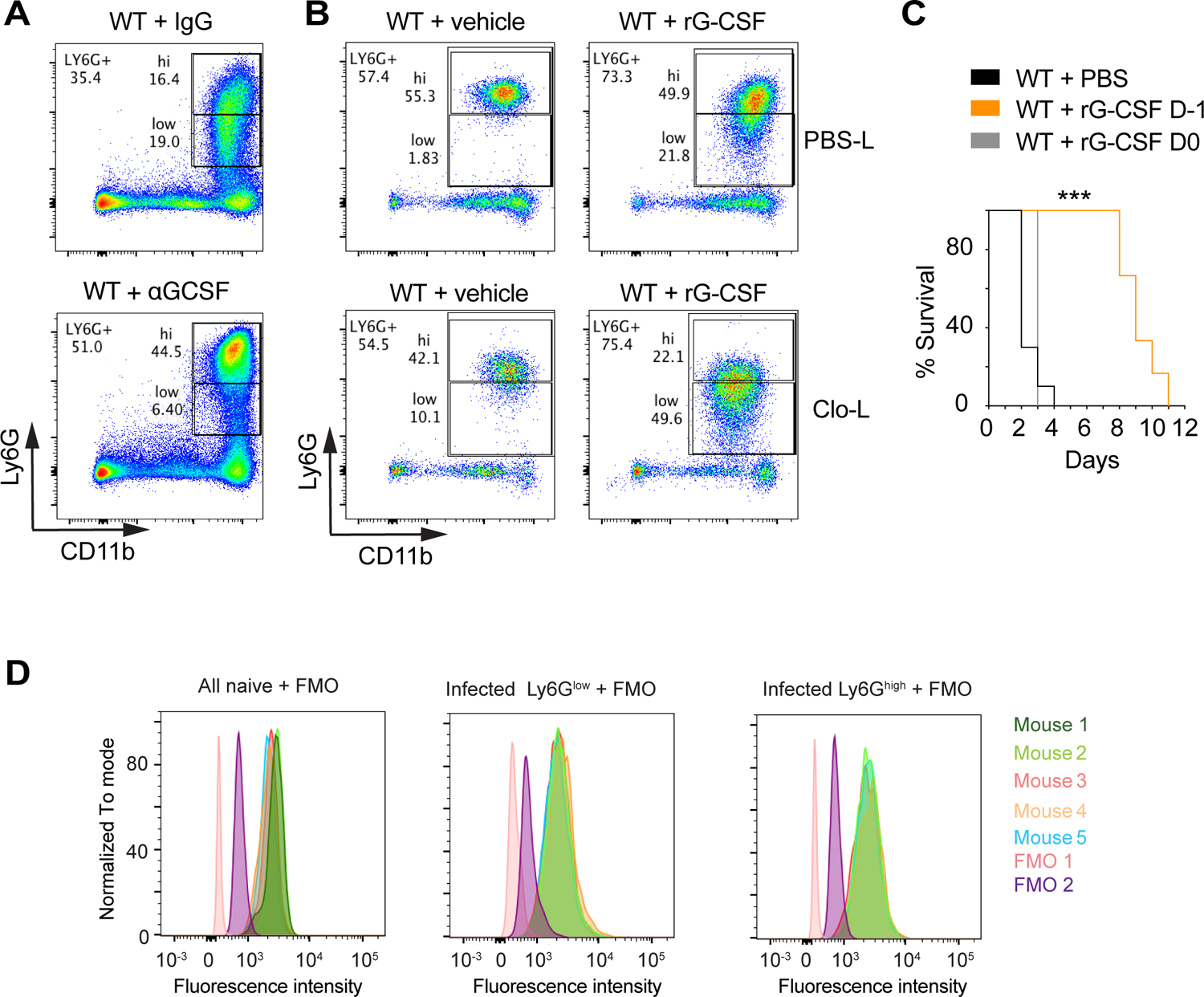
G-CSF and cell death-derived histones alter neutrophil populations. A. Representative flow cytometry plots assessing Ly6G^low^/Ly6G^high^ ratios in the spleens of WT mice infected with 5×10^5^ *C. albicans*, 72 hrs post-infection, treated with control or anti-G-CSF antibody at 24 hrs and 48 hrs post-infection. B. Representative flow cytometry plots of Ly6G^low^/Ly6G^high^ ratios of splenic neutrophils in mice pre-treated with PBS-L or Clo-L and injected with PBS or recombinant G-CSF (rG-CSF) for 24 hrs or 48 hrs and assessed 72 hrs after liposome administration. C. Survival of WT mice pre-treated with PBS of recombinant G-CSF, started either 24 hrs prior to infection or on the day of infection with 5×10^5^ WT *C. albicans*. D. Flow cytometric analysis of G-CSFR protein surface expression in Ly6G^high^ neutrophils from naïve WT mice (left panel), or Ly6G^high^ (middle panel) and Ly6G^low^ (right panel) neutrophils from infected mice 3-4 days post-infection. Controls represented by Ly6G^high^ neutrophils of naïve mice stained in the absence of specific primary- and secondary antibodies (FMO1) or in absence of primary but presence of secondary antibody (FMO2). Statistical analysis by Log-rank (Mantel-Cox) test for survival analysis (* p<0.05, ** p<0.01, *** p<0.001, **** p<0.0001)..

**Figure S8.**
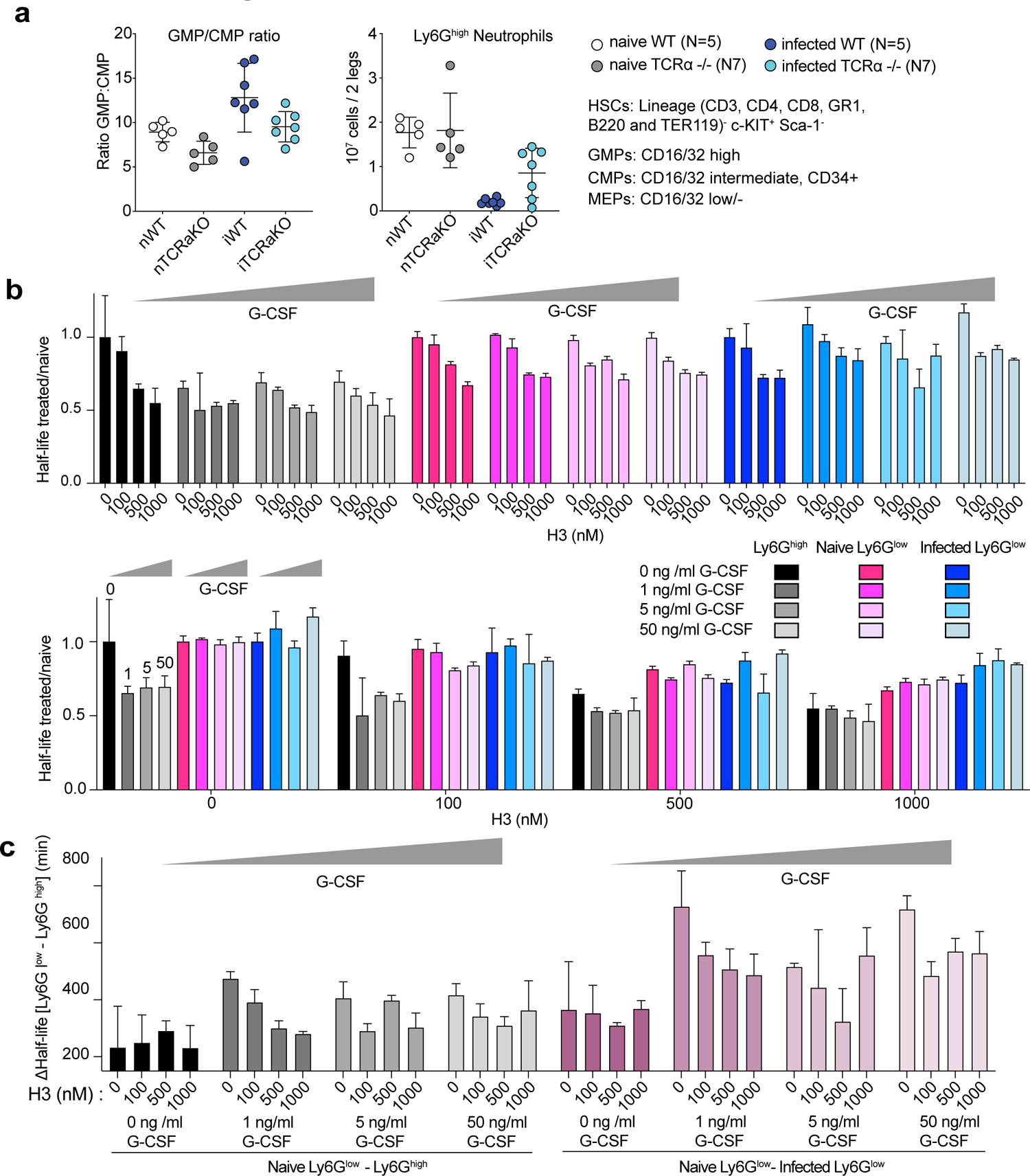
Effect of G-CSf and histones on neutrophil populations and their progenitors A. Flow cytometric analysis of myelopoietic progenitors and mature Ly6G^high^ neutrophils in the bone marrows of naive and infected WT and TCRα-deficient animals. In the far left panel, total common myeloid progenitors (CMP), megakaryocyte-erythrocyte progenitors (MEP) and granulocyte-monocyte progenitors (GMP) are shown. The middle panel depicts the ratios of GMPs to CMPs and the right panel the corresponding total Ly6G^high^ neutrophils in these bone marrows. B. Half-life ratio of Ly6G^high^ and Ly6G^low^ neutrophils from naive mice or Ly6G^low^ neutrophils from infected symptomatic mice supplemented with histone H3 or G-CSF alone or in combination at varying concentrations as indicated in the color scheme over naive (lane 1). Difference in half-life between WT naive Ly6G^high^ and Ly6G^low^ neutrophils (black and grey shades) or WT naive Ly6G^high^ and Ly6G^low^ Ly6G^low^ neutrophils from infected symptomatic mice (purple shades) alone or supplemented with histone H3 or G-CSF alone or in combination at the indicated concentrations.

## References

1. K. E. Rudd et al., Global, regional, and national sepsis incidence and mortality, 1990-2017: analysis for the Global Burden of Disease Study. Lancet 395, 200-211 (2020).

2. G. D. Brown et al., Hidden killers: human fungal infections. Sci Transl Med 4, 165rv113 (2012).

3. F. Bongomin, S. Gago, R. O. Oladele, D. W. Denning, Global and Multi-National Prevalence of Fungal Diseases-Estimate Precision. Journal of fungi 3, (2017).

4. B. G. Chousterman, F. K. Swirski, G. F. Weber, Cytokine storm and sepsis disease pathogenesis. Semin Immunopathol 39, 517–528 (2017).

5. J. Xu et al., Extracellular histones are major mediators of death in sepsis. Nat Med 15, 1318–1321 (2009).

6. J. Xu, X. Zhang, M. Monestier, N. L. Esmon, C. T. Esmon, Extracellular histones are mediators of death through TLR2 and TLR4 in mouse fatal liver injury. J Immunol 187, 2626–2631 (2011).

7. A. Havelka, K. Sejersen, P. Venge, K. Pauksens, A. Larsson, Calprotectin, a new biomarker for diagnosis of acute respiratory infections. Sci Rep 10, 4208 (2020).

8. H. Huang et al., Endogenous histones function as alarmins in sterile inflammatory liver injury through Toll-like receptor 9 in mice. Hepatology 54, 999–1008 (2011).

9. T. D. Tsourouktsoglou, et al., Histones, DNA, and Citrullination Promote Neutrophil Extracellular Trap Inflammation by Regulating the Localization and Activation of TLR4. Cell reports 31, 107602 (2020).

10. J. S. Boomer et al., Immunosuppression in patients who die of sepsis and multiple organ failure. Jama 306, 2594–2605 (2011).

11. S. C. Cheng et al., Broad defects in the energy metabolism of leukocytes underlie immunoparalysis in sepsis. Nat Immunol 17, 406–413 (2016).

12. J. J. Grailer, M. Kalbitz, F. S. Zetoune, P. A. Ward, Persistent neutrophil dysfunction and suppression of acute lung injury in mice following cecal ligation and puncture sepsis. J Innate Immun 6, 695–705 (2014).

13. F. Stephan et al., Impairment of polymorphonuclear neutrophil functions precedes nosocomial infections in critically ill patients. Critical care medicine 30, 315–322 (2002).

14. X. Xie et al., Single-cell transcriptome profiling reveals neutrophil heterogeneity in homeostasis and infection. Nat Immunol 21, 1119–1133 (2020).

15. R. Taneja, A. P. Sharma, M. B. Hallett, G. P. Findlay, M. R. Morris, Immature circulating neutrophils in sepsis have impaired phagocytosis and calcium signaling. Shock 30, 618–622 (2008).

16. R. S. Hotchkiss et al., Caspase inhibitors improve survival in sepsis: a critical role of the lymphocyte. Nat Immunol 1, 496–501 (2000).

17. *S.* *Roth* et al., Post-injury immunosuppression and secondary infections are caused by an AIM2 inflammasome-driven signaling cascade. Immunity, (2021).

18. Y. Zhang, et al., PD-L1 blockade improves survival in experimental sepsis by inhibiting lymphocyte apoptosis and reversing monocyte dysfunction. Critical care 14, R220 (2010).

19. B. Niu et al., Different Expression Characteristics of LAG3 and PD-1 in Sepsis and Their Synergistic Effect on T Cell Exhaustion: A New Strategy for Immune Checkpoint Blockade. Front Immunol 10, 1888 (2019).

20. X. Huang et al., PD-1 expression by macrophages plays a pathologic role in altering microbial clearance and the innate inflammatory response to sepsis. Proc Natl Acad Sci U S A 106, 6303–6308 (2009).

21. C. Nedeva et al., TREML4 receptor regulates inflammation and innate immune cell death during polymicrobial sepsis. Nat Immunol 21, 1585–1596 (2020).

22. F. Sonego et al., Paradoxical Roles of the Neutrophil in Sepsis: Protective and Deleterious. Front Immunol 7, 155 (2016).

23. Q. Guo et al., Induction of alarmin S100A8/A9 mediates activation of aberrant neutrophils in the pathogenesis of COVID-19. Cell Host Microbe 29, 222–235 e224 (2021).

24. S. J. Klebanoff, A. J. Kettle, H. Rosen, C. C. Winterbourn, W. M. Nauseef, Myeloperoxidase: a front-line defender against phagocytosed microorganisms. Journal of Leukocyte Biology 93, 185–198 (2013).

25. D. Kutter et al., Consequences of total and subtotal myeloperoxidase deficiency: risk or benefit ? Acta Haematol 104, 10–15 (2000).

26. K. Suzuki, H. Nunoi, M. Miyazaki, F. Koi, Prevalence of inherited myeloperoxidase deficiency in Japan. Peroxidase Multigene Family of Enzymes, 145-149 (2000).

27. K. D. Metzler, et al., Myeloperoxidase is required for neutrophil extracellular trap formation: implications for innate immunity. Blood 117, 953–959 (2011).

28. Y. Aratani et al., Differential host susceptibility to pulmonary infections with bacteria and fungi in mice deficient in myeloperoxidase. Journal of Infectious Diseases 182, 1276–1279 (2000).

29. N. Branzk et al., Neutrophils sense microbe size and selectively release neutrophil extracellular traps in response to large pathogens. Nat Immunol 15, 1017–1025 (2014).

30. K. D. Metzler, C. Goosmann, A. Lubojemska, A. Zychlinsky, V. Papayannopoulos, A myeloperoxidase-containing complex regulates neutrophil elastase release and actin dynamics during NETosis. Cell reports 8, 883–896 (2014).

31. J. S. Knight et al., Peptidylarginine deiminase inhibition reduces vascular damage and modulates innate immune responses in murine models of atherosclerosis. Circulation research 114, 947–956 (2014).

32. M. R. Rodrigues, D. Rodriguez, M. Russo, A. Campa, Macrophage activation includes high intracellular myeloperoxidase activity. Biochem Biophys Res Commun 292, 869–873 (2002).

33. K. Takahara, S. Tokieda, K. Nagaoka, K. Inaba, Efficient capture of Candida albicans and zymosan by SIGNR1 augments TLR2-dependent TNF-alpha production. Int Immunol 24, 89–96 (2012).

34. V. Papayannopoulos, D. Staab, A. Zychlinsky, Neutrophil Elastase Enhances Sputum Solubilization in Cystic Fibrosis Patients Receiving DNase Therapy. Plos One 6, (2011).

35. S. Basu et al., “Emergency” granulopoiesis in G-CSF-deficient mice in response to Candida albicans infection. Blood 95, 3725–3733 (2000).

36. B. E. Hsu et al., Immature Low-Density Neutrophils Exhibit Metabolic Flexibility that Facilitates Breast Cancer Liver Metastasis. Cell reports 27, 3902–3915 e3906 (2019).

37. H. Knaup et al., Early therapeutic plasma exchange in septic shock: a prospective open-label nonrandomized pilot study focusing on safety, hemodynamics, vascular barrier function, and biologic markers. Critical care 22, 285 (2018).

38. X. Cheng, J. E. Ferrell, Jr., Apoptosis propagates through the cytoplasm as trigger waves. Science 361, 607–612 (2018).

39. C. Dubois et al., High plasma level of S100A8/S100A9 and S100A12 at admission indicates a higher risk of death in septic shock patients. Sci Rep 9, 15660 (2019).

40. A. Larsson et al., Calprotectin is superior to procalcitonin as a sepsis marker and predictor of 30-day mortality in intensive care patients. Scand J Clin Lab Invest 80, 156–161 (2020).

41. J. Dominguez-Andres et al., Inflammatory Ly6C(high) Monocytes Protect against Candidiasis through IL-15-Driven NK Cell/Neutrophil Activation. Immunity 46, 1059–1072 e1054 (2017).

42. P. J. Jenks, E. Jones, Infections in asplenic patients. Clinical microbiology and infection: the official publication of the European Society of Clinical Microbiology and Infectious Diseases 1, 266–272 (1996).

43. Y. S. Kang et al., The C-type lectin SIGN-R1 mediates uptake of the capsular polysaccharide of Streptococcus pneumoniae in the marginal zone of mouse spleen. Proc Natl Acad Sci U S A 101, 215–220 (2004).

44. A. Lanoue et al., SIGN-R1 contributes to protection against lethal pneumococcal infection in mice. J Exp Med 200, 1383–1393 (2004).

45. P. D. Uchil et al., A Protective Role for the Lectin CD169/Siglec-1 against a Pathogenic Murine Retrovirus. Cell Host Microbe 25, 87–100 e110 (2019).

46. X. Sewald et al., Retroviruses use CD169-mediated trans-infection of permissive lymphocytes to establish infection. Science 350, 563–567 (2015).

47. J. P. Gaut et al., Neutrophils employ the myeloperoxidase system to generate antimicrobial brominating and chlorinating oxidants during sepsis. Proc Natl Acad Sci U S A 98, 11961–11966 (2001).

48. S. Hamaguchi et al., Origin of Circulating Free DNA in Sepsis: Analysis of the CLP Mouse Model. Mediators Inflamm 2015, 614518 (2015).

49. M. A. Stark et al., Phagocytosis of apoptotic neutrophils regulates granulopoiesis via IL-23 and IL-17. Immunity 22, 285–294 (2005).

50. D. Zhang et al., Neutrophil ageing is regulated by the microbiome. Nature 525, 528–532 (2015).

51. J. M. Adrover et al., Programmed ‘disarming’ of the neutrophil proteome reduces the magnitude of inflammation. Nat Immunol 21, 135–144 (2020).

52. J. M. Adrover et al., A Neutrophil Timer Coordinates Immune Defense and Vascular Protection. Immunity 50, 390–402 e310 (2019).

53. M. Casanova-Acebes et al., Rhythmic modulation of the hematopoietic niche through neutrophil clearance. Cell 153, 1025–1035 (2013).

54. H. J. Kwak et al., Myeloid cell-derived reactive oxygen species externally regulate the proliferation of myeloid progenitors in emergency granulopoiesis. Immunity 42, 159–171 (2015).

55. J. J. Lai, F. M. Cruz, K. L. Rock, Immune Sensing of Cell Death through Recognition of Histone Sequences by C-Type Lectin-Receptor-2d Causes Inflammation and Tissue Injury. Immunity 52, 123–135 e126 (2020).

56. G. D. Brown, Dectin-1: a signalling non-TLR pattern-recognition receptor. Nature reviews. Immunology 6, 33–43 (2006).

57. A. Warnatsch et al., Reactive Oxygen Species Localization Programs Inflammation to Clear Microbes of Different Size. Immunity 46, 421–432 (2017).

58. M. Demers et al., Priming of neutrophils toward NETosis promotes tumor growth. Oncoimmunology 5, e1134073 (2016).

59. M. Demers et al., Cancers predispose neutrophils to release extracellular DNA traps that contribute to cancer-associated thrombosis. Proc Natl Acad Sci U S A 109, 13076–13081 (2012).

60. B. M. Szczerba et al., Neutrophils escort circulating tumour cells to enable cell cycle progression. Nature 566, 553–557 (2019).

61. L. Yang et al., DNA of neutrophil extracellular traps promotes cancer metastasis via CCDC25. Nature 583, 133–138 (2020).

62. F. Wen, A. Shen, A. Choi, E. W. Gerner, J. Shi, Extracellular DNA in pancreatic cancer promotes cell invasion and metastasis. Cancer Res 73, 4256–4266 (2013).

63. J. Park et al., Cancer cells induce metastasis-supporting neutrophil extracellular DNA traps. Sci Transl Med 8, 361ra138 (2016).

64. J. Schulte-Schrepping et al., Severe COVID-19 Is Marked by a Dysregulated Myeloid Cell Compartment. Cell 182, 1419–1440 e1423 (2020).

65. Y. Zuo, et al., Neutrophil extracellular traps in COVID-19. JCI Insight 5, (2020).

66. C. B. Messner et al., Ultra-High-Throughput Clinical Proteomics Reveals Classifiers of COVID-19 Infection. Cell Syst 11, 11–24 e14 (2020).

67. V. Demichev, C. B. Messner, S. I. Vernardis, K. S. Lilley, M. Ralser, DIA-NN: neural networks and interference correction enable deep proteome coverage in high throughput. Nat Methods 17, 41–44 (2020).

68. A. Dobin et al., STAR: ultrafast universal RNA-seq aligner. Bioinformatics 29, 15–21 (2013).

69. B. Li, C. N. Dewey, RSEM: accurate transcript quantification from RNA-Seq data with or without a reference genome. BMC Bioinformatics 12, 323 (2011).

70. D. S. DeLuca et al., RNA-SeQC: RNA-seq metrics for quality control and process optimization. Bioinformatics 28, 1530–1532 (2012).

71. M. I. Love, W. Huber, S. Anders, Moderated estimation of fold change and dispersion for RNA-seq data with DESeq2. Genome Biol 15, 550 (2014).

72. A. Subramanian et al., Gene set enrichment analysis: a knowledge-based approach for interpreting genome-wide expression profiles. Proc Natl Acad Sci U S A 102, 15545–15550 (2005).

